# Neutrophil-microglia interaction drives reversible motor dysfunction in neuromyelitis optica model induced by subarachnoid AQP4-IgG

**DOI:** 10.1101/2025.08.22.671883

**Authors:** Fangfang Qi, Vanda A. Lennon, Shunyi Zhao, Yong Guo, Husheng Ding, Caiyun Liu, Whitney M. Bartley, Tingjun Chen, Claudia F. Lucchinetti, Long-Jun Wu

## Abstract

Neutrophils and neutrophil extracellular traps (NETs) contribute to early neuromyelitis optica (NMO) histopathology initiated by IgG targeting astrocytic aquaporin-4 water (AQP4) channels. Yet, the mechanisms recruiting neutrophils and their pathogenic roles in disease progression remain unclear. To investigate molecular-cellular events preceding classical complement cascade activation in a mouse NMO model, we continuously infused, via spinal subarachnoid route, a non-complement-activating monoclonal AQP4-IgG. Parenchymal infiltration of netting neutrophils containing C5a ensued with microglial activation and motor impairment, but no blood–brain barrier leakage. Motor impairment and neuronal dysfunction both reversed when AQP4-IgG infusion stopped. Two-photon microscopy and electron-microscopy-based reconstructions revealed physical interaction of infiltrating neutrophils with microglia. Ablation of either peripheral neutrophils or microglia attenuated the motor deficit, highlighting their synergistic pathogenic roles. Of note, mice lacking complement receptor C5aR1 exhibited reduction in neutrophil infiltration, microglial lysosomal activation, neuronal lipid-droplet burden and motor impairment. Pharmacological inhibition of C5aR1 recapitulated this protection. Immunohistochemical analysis of an NMO patient’s early spinal cord lesions revealed analogous pathological findings. Our study identifies neutrophil-derived C5a signaling through microglial C5aR1 as a key early driver of reversible motor neuron dysfunction in the precytolytic phase of NMO.

**One Sentence Summary:** Neutrophil-derived C5a coactivates microglia to drive reversible motor paresis initiated by a non-complement-activating aquaporin-4-IgG binding to astrocytes in a mouse model of neuromyelitis optica.

## INTRODUCTION

Neuromyelitis optica (NMO) spectrum disorder is a severe, relapsing, inflammatory, autoimmune disorder of the central nervous system (CNS) with myelin loss occurring secondary to astrocyte targeting by a complement-activating IgG specific for the aquaporin-4 (AQP4) water channel ^1–3^. A C5-neutralizing-IgG (eculizumab) is highly effective therapeutically in terminating acute NMO attacks and preventing relapse ^4^. Untreated, the fully established NMO lesion shows extensive deposition of the C5b-C9 membrane attack complex on astrocytes, blood–brain barrier (BBB) leakage and gross tissue destruction ^5^. These late events in the acute attack phase require substantial BBB disruption to permit CNS influx of plasma IgG, rheumatoid-factor-like IgM ^6^ and liver-derived complement components.

The presence of neutrophils in early CNS lesions ^5,7,8^ is a histopathological feature distinguishing NMO from other inflammatory immune-mediated demyelinating diseases, such as multiple sclerosis which generally lack neutrophils ^9,10^. Neutrophil invasion of the spinal cord ^5^ and brain parenchyma ^7^ is thought to contribute to astrocyte dysfunction and tissue damage ^11^.

Microglia, as CNS immune sentinel cells, continuously monitor and respond to parenchymal changes ^12–15^. A mouse model of NMO established in this laboratory demonstrated that motor neuronal dysfunction requires microglial activation by complement component C3a emanating from AQP4-IgG-activated astrocytes ^16^. However, it remains unknown how resident microglia interact with infiltrating neutrophils and influence NMO lesion progression. To investigate early pathogenic events preceding classical complement cascade activation in evolving NMO lesions, we modified our established mouse model ^16^ by infusing continuously, via spinal subarachnoid route, a non-complement-activating IgG specific for the AQP4 extracellular domain. Hypothesizing a pathophysiological role for CNS-infiltrating neutrophils in initiating early NMO lesions through chemotaxis ^11^ and cytokine release ^17,18^, we employed high-parameter flow cytometry and electron microscopy to characterize these neutrophils and their interactions. Our findings reveal that physical neutrophil-microglia interaction is a pre-requisite for neurological impairment in the precytolytic phase of NMO. Early NMO deficits that in patients are sometimes reversible by plasma exchange and anti-inflammatory corticosteroid therapy ^19^ are driven in the mouse model by neutrophil-derived C5a. Genetic deletion of C5a receptor 1 (C5aR1) significantly reduced neutrophil infiltration, motor neuron lipid-oxidative stress, and functional motor impairment.

## RESULTS

### Neutrophils infiltrate the spinal parenchyma early in the mouse model of NMO

We initially reported microglial activation and motor impairment as outcomes of 5 days’ continuous lumbar subarachnoid infusion of complement-activating serum IgG derived from NMO patients or a complement-activating monoclonal mouse AQP4-IgG ^16^. To investigate the contribution of neutrophils in the pre-cytolytic stage of NMO, we infused a non-complement-activating mouse monoclonal IgG1 specific for the AQP4 extracellular domain (Figure 1A). High-spectrum flow cytometry revealed time-dependent infiltration of neutrophils into spinal parenchyma. Numbers peaked on day 3, with CD11b^+^Ly6G^+^Ly6C^neg/low^ neutrophils accounting for 5.1% of CD45^+^ immune cells; on day 5, 3.9% were neutrophils (Figure 1B and Figure S1). Lumbar spinal cord immunostaining on day 3 confirmed neutrophils infiltrating the parenchyma and revealed a greater abundance of neutrophils with extracellular traps (NETs, myeloperoxidase [MPO]-positive; Figure 1C-E) as well as Ly6G-positive neutrophils (Figure 1 F and G) in ventral cord grey matter, partly surrounding a blood vessel-like structure (Figure 1, *D1* and rectangle F1 in F). The images of neutrophils within or near blood vessels in the spinal parenchyma of AQP4-IgG recipient mice are consistent with transvascular infiltration (Figure 1 H). On day 3, neutrophil elastase (NE)-positive neutrophils were scattered in the ventral horn, dorsal horn and meninges (Figure 1 I); infiltrating NET^+^-neutrophils were 5-6-fold more numerous in AQP4-IgG recipients than in control-IgG recipients (Figure 1 E, G and I). Our results indicate that AQP4-IgG infused into the lumbar subarachnoid space of mice triggers neutrophil migration into the neighboring cord parenchyma, with a subset undergoing NETosis. Thus, our model closely resembles early-stage lesions observed in NMO patients ^8^.

**Figure 1.**
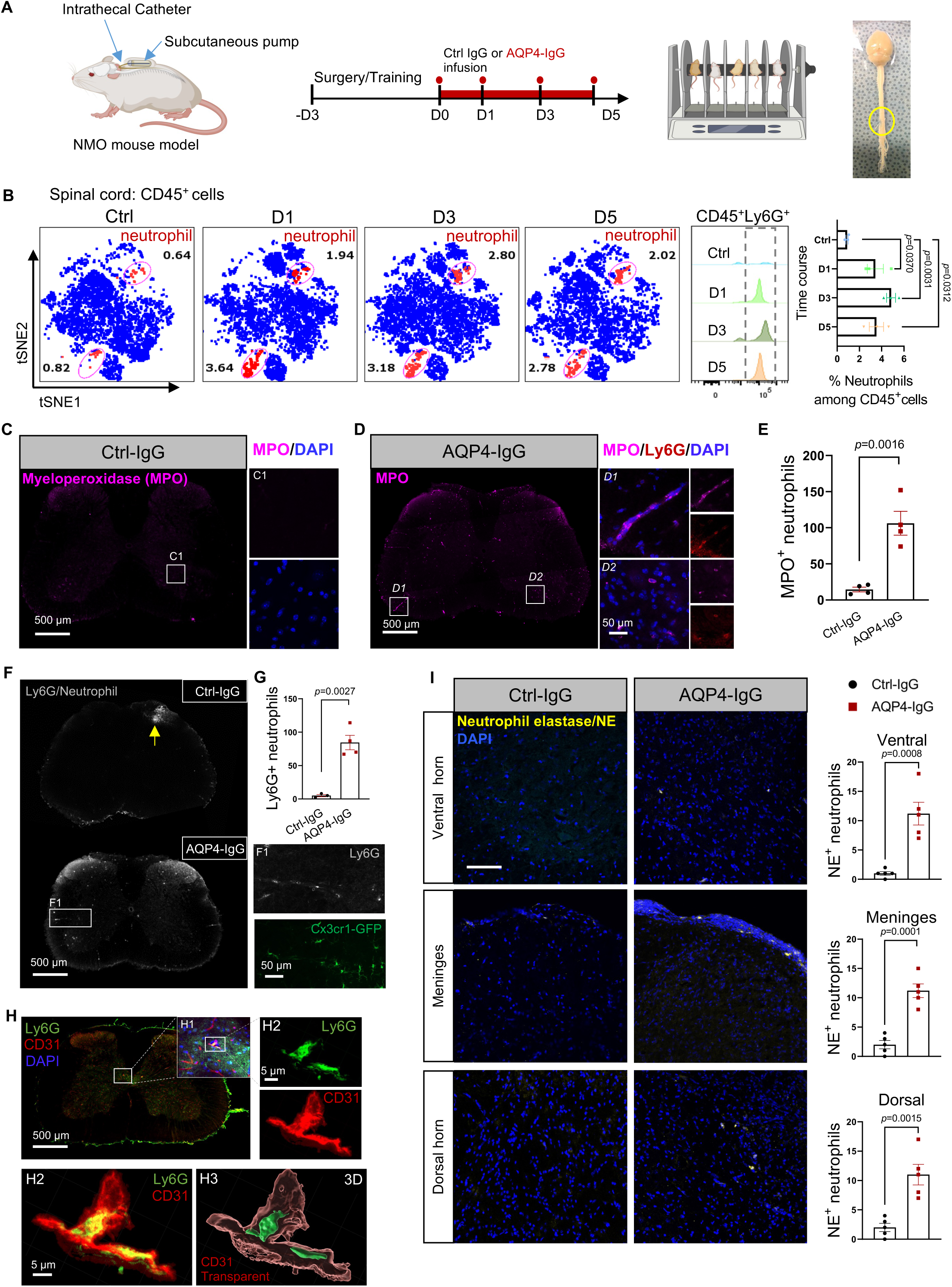
Neutrophil infiltration and NETosis in the spinal cord of a mouse model of neuromyelitis optica. A. Experimental design: Catheter is inserted surgically via cisterna magna into L4 subarachnoid space (yellow circle, verified by Evans blue infusion). Osmotic pump, placed subcutaneously day 0, continuously infuses mouse monoclonal AQP4-IgG (AQP4-ECD-m21-IgG1) or control mouse IgG (1.2 μg/day, in 12 µL). Rotarod motor training, days –3, –2, –1; testing, days 0, +1, +2, +3, +4 and +5. Terminal transcardiac perfusion: spinal cord tissue harvested for immunohistochemical and flow cytometric analyses. B. Cytek analysis of spinal cord-infiltrating neutrophils; days 1, 3, and 5 of IgG infusion. Control mice (1^st^ panel) were infused 3 days with normal mouse IgG. Mice in panels 2-4 received AQP4-IgG. *n* = 3 mice per group. C-E. Representative confocal images at day 3 of IgG infusion (C, D) and quantification (E) of neutrophils (Ly6G^+^/ MPO^+^) in lumbar cord parenchyma (*n*= 4 mice per group). C1. Higher magnification of box in C shows neutrophils (magenta) and cell nuclei (blue). *D1*, *D2*. Higher magnification of Ly6G^+^/ MPO^+^ netting neutrophils in AQP4-IgG-infused cord. F, G. Representative confocal images at day 3 of IgG infusion. Lumbar cord of Ctrl-IgG and AQP4-IgG recipients; Ly6G^+^ cluster at top right of control dorsal cord (yellow arrow) is subarachnoid inflammation at catheter site. Higher magnification images (F1), at right of boxed area, reveal neutrophils in white matter, in or around a blood vessel of AQP4-IgG recipients; Cx3cr1-GFP^+^ microglia/macrophage surround the blood vessel (F). Quantification of Ly6G^+^ neutrophils in cord section; *n* = 4 mice per group (G). H. Representative confocal images of CD31^+^ blood vessels and Ly6G^+^ neutrophils at day 3 in lumbar cord of AQP4-IgG recipient mouse. H1. Higher magnification of boxed Ly6G^+^/CD31^+^ in H. H2. Higher magnification (split and merged) of boxed Ly6G^+^ and CD31^+^ in H1. H3. 3D Imaris rendering of Ly6G^+^ neutrophils (green) and CD31^+^ blood vessel (red; transparent) in H2. I. Lumbar cord distribution of NE^+^ neutrophils in ventral horn, meninges, and dorsal horn at day 5 of control-IgG or AQP4-IgG infusion. *n* = 4 mice per group. Unpaired Student *t*-test. All statistical tests shown for quantitative data (means ± SEM) in graphs of Figure 1 represent two-sided statistical tests. One-way ANOVA, followed by Tukey *post hoc* multiple comparisons test (B). Unpaired Student *t*-test in E, G and I. *p* < 0.05 was considered a significant difference. Exact *p* values and scales are in the figure panels.

### Neutrophils interact with microglia in NMO

Microglia potentially respond to local cues from many cell types in evolving CNS lesions induced by subarachnoid AQP4-IgG infusion ^16,20–22^. In mice with genetically GFP-tagged microglia (Cx3cr1GFP), confocal images of lumbar cord revealed Ly6G-immunostained neutrophils in proximity to microglia (Figure 2A). Imaris 3D rendering of those images revealed potential foci of neutrophil-microglia physical interaction (Figure 2A1). Near-infrared branding by 2-photon microscopy (Figure 2B) allowed electron microscopic (EM) acquisition of higher resolution images (Figure 2C and Figure S2A-C). Serial block-face scanning electron microscopy enabled the cells of interest observed in confocal images to be identified: microglia (green), neutrophil (red), and two neurons (blue) (Figure 2C and Movie S1). 3D reconstruction based on serial EM images and confocal images confirmed two types of physical neutrophil-microglial interactions (Figure 2C1, 2C2, 2D and 2E): neutrophil soma-microglial process and neutrophil soma-microglial soma. The percentage of each neutrophil interaction type was enumerated by employing high resolution Z-stacked confocal images of Ly6G and MPO immunoreactivities and subsequent 3D rendering (Figure 2E, 2F, and Movie S2-1, 2-2, and 3): 47% contacted microglial processes (Figure 2E1 and 2F) and 19% contacted microglial somata (Figure 2E2); approximately 34% had no microglial contact (Figure S2D).

**Figure 2.**
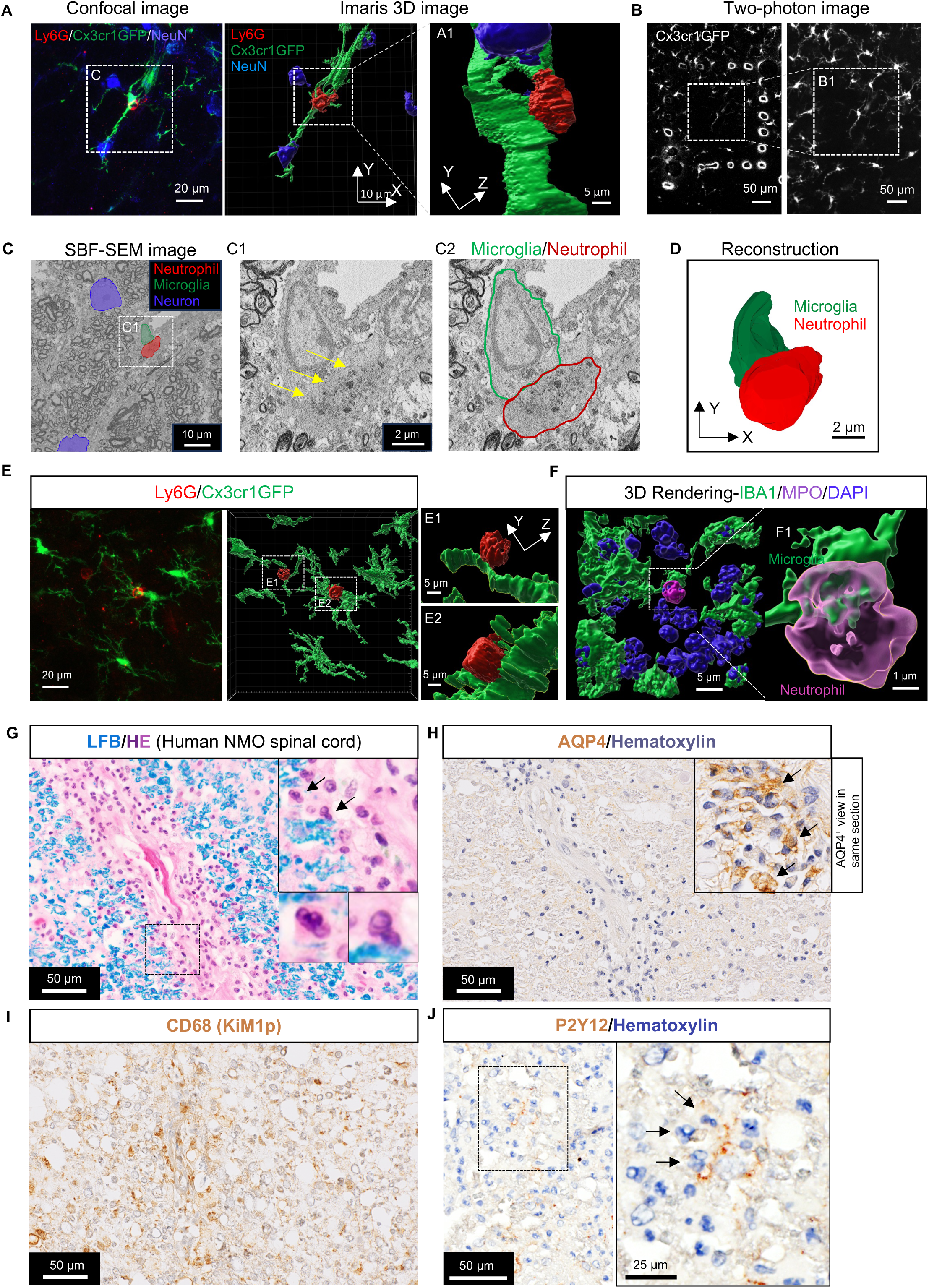
Neutrophil-microglial contacts in lumbar parenchyma at day 3 of AQP4-IgG infusion. A. Confocal image identifies putatively interacting microglial cell (Cx3cr1GFP^+^, green) and neutrophil (Ly6G^+^, red) adjacent to a neuronal soma (NeuN^+^ nucleus, blue); subsequent images with Imaris 3D rendering (A1). B. Laser branding under two-photon imaging frames the cord region of interest containing the contacting neutrophil-microglia; magnified in B1. C. Serial block-face scanning electron microscopy (SBF-SEM) of the region of interest shows, at the ultrastructural level, the same neutrophil, microglia and 2 neurons. Yellow arrows in magnified C1 box indicate contacting edges of microglial and neutrophil somata (outlined by green and red lines in C2). D. 3D serial reconstruction of contacting microglial and neutrophil somata shown in C (See Video S1, Z-stack). E. Representative confocal image (left, 2,048 × 2,048 pixel; ×63 objective lens) and 3D rendering image (right) show an interacting microglial process (E1) and soma (E2) with Ly6G^+^ neutrophils in lumbar cord parenchyma of an AQP4-IgG infused mouse (see Videos S2-1, S2-2, Imaris). F. Representative Imaris 3D rendering image shows an (MPO^+^) neutrophil undergoing netting and interacting with microglial processes in lumbar cord parenchyma of an AQP4-IgG-infused mouse. F1. High power shows microglial process physically interacting with neutrophil soma (see video S3, Imaris). G-I. Serial histopathologic sections of an NMO patient early spinal cord lesion. G. White matter parenchyma shows intact myelin (Luxol fast blue) with neutrophils and eosinophils infiltrating the parenchyma in close proximity to a penetrating blood vessel. The right upper inset shows the area of neutrophil infiltration, the right lower inset shows two infiltrated neutrophils (arrow identifies their multi-lobed nuclei and light pink eosinophilic cytoplasm). H. Immunohistochemistry reveals reduction of AQP4 immunoreactivity in this region. Inset at right shows AQP4 retention in adjacent area I. Macrophages/microglia identified by CD68 (KiM1P) immunostain in this non-demyelinated lesion. J. Neutrophils among P2Y12 positive microglia in this non-demyelinated lesion. Magnified view (inset box right) shows four neutrophils (segmented nuclei, hematoxylin-stained) in close proximity to microglial processes.

To investigate the pertinence of these findings to lesion evolution in NMO patients, we analyzed immunohistochemically early lesions in the spinal cord of an NMO patient. In white matter (Figure 2G), myelin remained intact and a penetrating blood vessel was surrounded by abundant granulocytes (predominantly neutrophils). AQP4 immunoreactivity in that region was reduced (Figure 2H) by comparison with an adjacent region in the same section (inset), and microglial/macrophage activation was prominent (Figure 2I). Consistent with findings in the mouse model, we observed P2Y12 microglia interacting with neutrophils in early pre-demyelinating lesions within this patient’s spinal white matter (Figure 2J). We next assessed blood-brain barrier integrity in the mouse model by evaluating, on days 3 and 5 of AQP4-IgG infusion, the distribution of endogenous fibrinogen (340 kDa) and dextran red (70 kDa) injected 40 minutes prior to perfusion (Figure S6A-C) and of the immunostained endothelial tight junction protein, Claudin-5. No BBB leakage was evident (Figure S6D). Thus, the prominent neutrophil infiltration is not due to BBB compromise. Documentation of CNS-infiltrating neutrophils interacting physically with microglia in early spinal cord lesions of AQP4-IgG-infused mice and in a patient with acute NMO supports a pathogenic role for neutrophils in early-stage NMO lesions.

### Neutrophils drive microglial activation and motor impairment initiated by AQP4-IgG binding to astrocytes

To investigate potential roles of infiltrating neutrophils in the neuropathology and motor dysfunction induced by AQP4-IgG, we depleted neutrophils by twice injecting Ly6G-specific IgG (100 mg/kg *i.p.*), 2 days before subarachnoid catheter insertion (Figure 3A and Figure S3). Flow cytometry confirmed loss of circulating neutrophils (Figure 3B); the percentage of neutrophils (CD45^+^CD11b^+^Ly6G^+^Ly6C^low^) among total circulating immune cells (CD45^+^CD11b^+^) was significantly lower in Ly6G-IgG-treated mice than in isotype control IgG-treated mice (t-SNE heatmap, Figure 3C). Similarly, peripheral MPO^+^ (NET^+^) cells were rare (see sectioned lung tissue of Ly6G-IgG-treated mice, Figure 3D). Mice treated with isotype control IgG exhibited worsening motor impairment and morphologic evidence of microglial activation in lumbar cord parenchyma during AQP4-IgG infusion; neither outcome was observed in neutrophil-depleted recipients of AQP4-IgG (Figure 3E, F, G and Movie S4). The peak of neutrophil infiltration (day 3 of AQP4-IgG infusion) coincided with morphologically evident microglial activation but preceded peak motor impairment (Figure 3G) which continued beyond day 5 but reversed spontaneously over the course of 3 weeks after AQP4-IgG infusion was stopped at day 7 (Figure 3H and I). The significant correlation between Rotarod latency to fall and microglial area expansion (Figure 3J) suggests that microglial activity influences the severity of motor impairment. Indeed, microglial ablation did mitigate motor impairment (Figure S4), consistent with our earlier report ^16^. These data indicate that, in NMO lesion evolution, infiltrating neutrophils or their secreted products enhance microglial activation and motor impairment initiated by AQP4-IgG binding to astrocytes.

**Figure 3.**
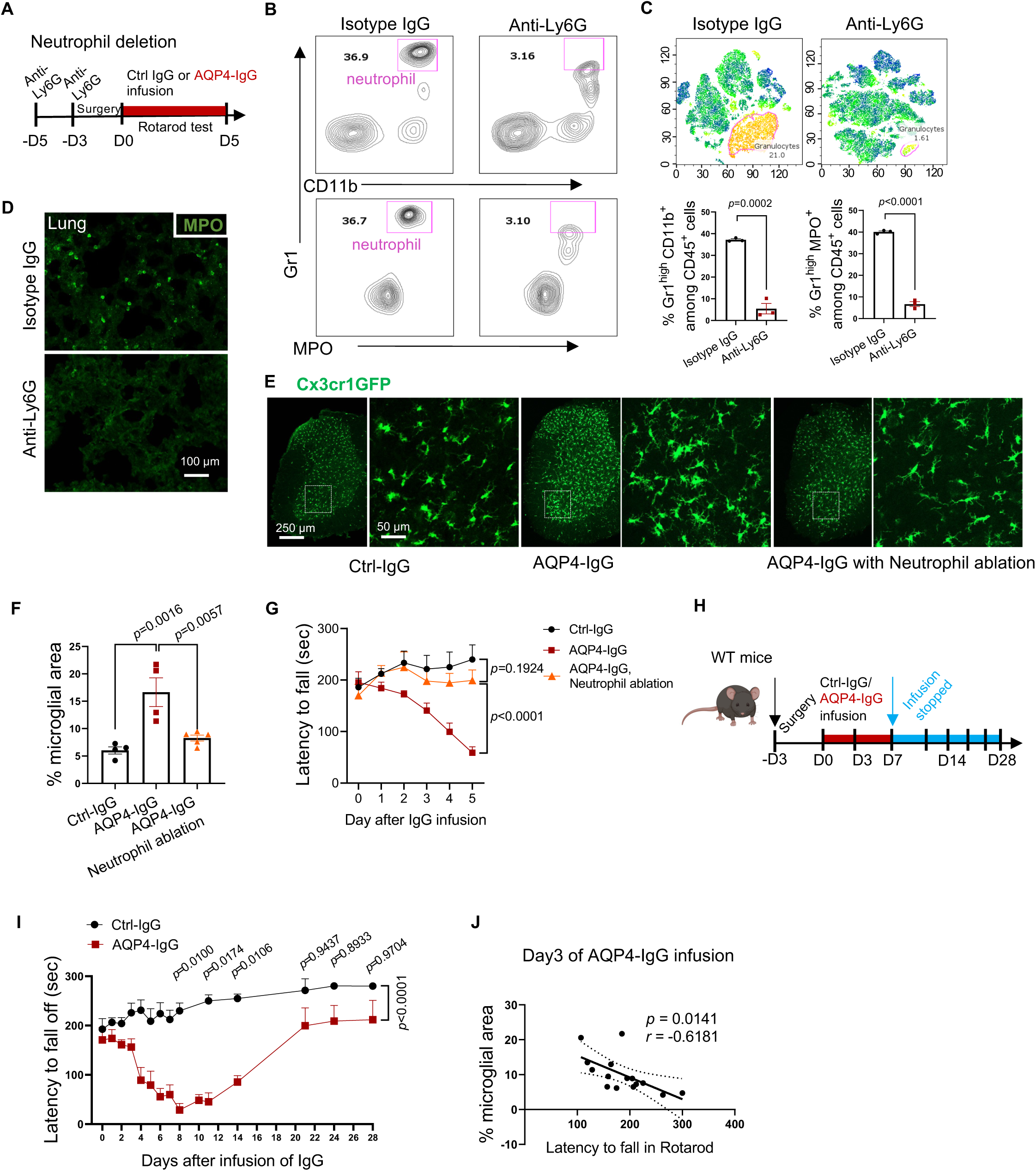
Microglial activation and motor impairment by AQP4-IgG requires CNS-infiltrating neutrophils. A. Timeline for *i.p.* injections of neutrophil-depleting anti-Ly6G-IgG, or isotype control IgG (100 mg/kg), lumbar subarachnoid catheter insertion, AQP4-IgG infusion and Rotarod testing. B. Flow cytometric analysis confirms neutrophil ablation efficiency (percentage CD45^+^CD11b^+^Gr1^+^MPO^+^ cells among peripheral CD45^+^CD11b^+^ cells). C. (upper) t-SNE analysis of CD45^+^ immune cell subtypes released from enzyme-dissociated lumbar spinal cord tissues of control and neutrophil-depleted mice. (lower) Quantification of data in B (*n* = 3 mice per group). D. Representative confocal images of peripheral neutrophils in lungs of mice receiving neutrophil-depleting anti-Ly6G IgG or isotype control IgG (*n* = 3 mice per group). E. Microglial activation, reflected by Cx3cr1GFP signal, in corresponding regions of lumbar cord of mice without and with neutrophil ablation (by Ly6G-IgG or isotype control IgG) after 3-days’ infusion with normal control mouse IgG or AQP4-IgG. F. Quantification of microglia-occupied areas in E (*n* = 4-5 mice per group). G. Motor function, reflected by fall latency from rotarod, in neutropenic mice (anti-Ly6G IgG-treated) and non-neutrophil-ablated (isotype control IgG-treated) during 5-days’ infusion of AQP4-IgG or normal control mouse IgG (0.1 µg/µL; time: *F* _(2.415, 28.98)_ = 4.838, *p*=0.0113; treatment:; *F*_(2, 12)_ = 12.46, *p*=0.0012; interaction: *F* _(10, 60)_ = 7.100, *p* < 0.0001; *n* = 5 mice per group). H. Experimental design: WT mice were continuously infused with Ctrl-IgG or AQP4-IgG by osmotic pumps for 7 days from day 0; infusion was discontinued at day 8. I. Motor impairment worsened progressively in AQP4-IgG recipients, with nadir at day 8. Continued rotarod testing for another 3 weeks showed progressive motor recovery from day 8. J. Correlations between microglial activation state (lumbar microglial area) and latency to fall in rotarod test. Simple linear regression (one dot represents one mouse at day 3 of IgG infusion). In all graphs, data represent means ± SEM and all statistical tests are two-sided. Unpaired Student *t*-test in C. One-way ANOVA, followed by Tukey *post hoc* multiple comparisons test in F. Two-way repeated measures ANOVA with Sidak *post hoc* test in G, I. *p* < 0.05 was considered a significant difference.

### Neutrophil-derived C5a enhances NETosis and microglial activation to initiate motor impairment

Levels of C5a, the chemoattractant polypeptide cleavage product of complement C5 protein that recruits and activates neutrophils ^23,24^, are elevated in CSF of patients with established NMO, in both attack and remission stages ^25^. We initially assumed that the source of C5a attracting early CNS infiltration of neutrophils was C5 protein produced and secreted by AQP4-IgG-activated astrocytes ^26^. However, we found no C5 proenzyme immunoreactivity in any spinal parenchymal cell of mice infused 5 days with AQP4-IgG (Figure S5). Further, we found no C5a immunoreactivity in astrocytes (Figure 4A), and very few IBA1^+^ macrophages/microglia or infiltrating CCR2^+^ monocytes were C5a-positive (Figure 4B and C). Instead, the cytoplasm of infiltrating MPO^+^ neutrophils accounted for >90% of C5a immunoreactivity (Figure 4C-E). Thus, in the evolving spinal cord lesion of this mouse NMO model, neutrophils appear to be the initial source of C5a. We next evaluated C5a receptor 1 (C5aR1) expression on subsets of leukocytes enzymatically dissociated from spinal cord tissue of IgG-infused mice (Figure 4F; gating strategy; Figure S1). Microglia and neutrophils highly expressed C5aR1; monocytes and lymphocytes expressed low levels (Figure 4 G). These data accord with a public transcriptomic database that showing high *C5aR1* mRNA expression in microglia/macrophages of non-perturbed mouse CNS tissue (Figure 4 H, *Brain RNA-Seq*). The scarcity of lumbar cord-infiltrating neutrophils on day 3 in *C5aR1*-null mice (Figure 4I and J) implicates C5a-C5aR1 signaling as the mediator of neutrophil infiltration and subsequent neutrophil-microglial interaction. Day 3 is the peak of neutrophil infiltration in wild-type recipients of AQP4-IgG. Consistent with our observations in neutrophil-immunodepleted mice (Figure 3G), genetic deletion of *C5aR1* or pharmacological inhibition of its function by PMX 205 (10 mg/kg) (Figure 4K and L) mitigated motor impairment.

**Figure 4.**
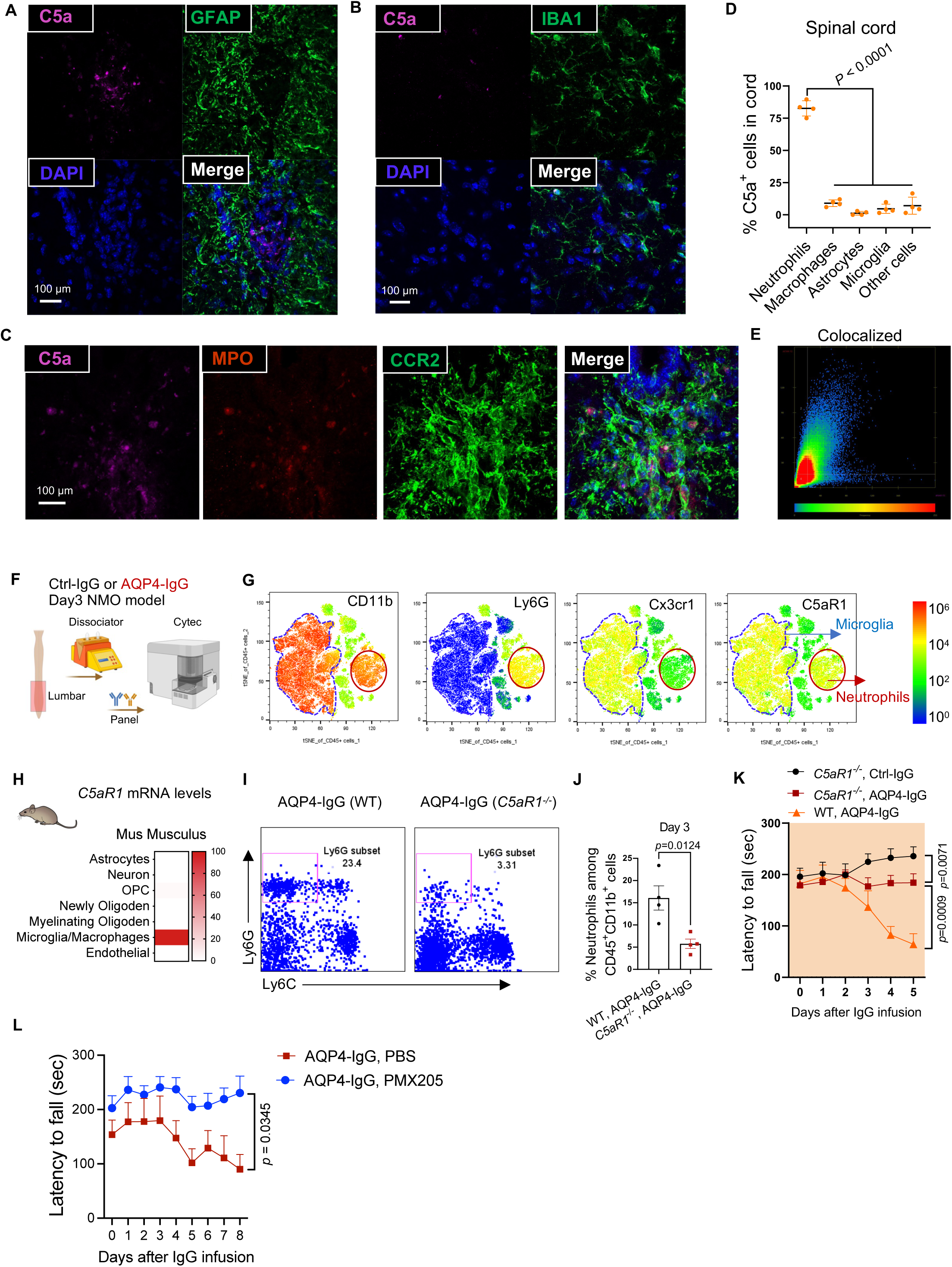
Neutrophil-derived C5a enhances NET production and activates microglia via C5aR1 signaling. A and B. In the spinal grey matter of AQP4-IgG-recipient mice, C5a-immunoreactivity was not seen in GFAP^+^ astrocytes (A) and rarely seen in IBA1^+^ microglia/macrophage (B). The C5 proenzyme was seen only in luminal plasma, not in any CNS parenchymal cells (Figure S4B). C. Triple immunostaining of lumbar cord of AQP4-IgG recipient mice revealed C5a in neutrophils (MPO+, left) and monocytes (CCR2-GFP^+^, enhanced by anti-GFP-IgG, right). D. The percentage of spinal cord cells expressing C5a immunoreactivity in AQP4-IgG-infused mice (determined by Image J) (*n*=4 mice in each group). E. Colocalization analysis of C5a and MPO signals in C using Zen software (LSM980). F. After transcardiac washout of vasculature, lumbar cords from IgG-infused mice were dissociated enzymatically and subjected to high parametric flow cytometric analysis (Cytek). G. tSNE maps identifying CD11b^+^, Ly6G^+^, Cx3cr1^+^ and C5aR1^+^ cells and their expression levels among CD45^+^ immune cells in the spinal cord, revealed that C5aR1 is expressed predominantly in microglial and neutrophil subsets. H. Public database (*Brain RNA-Seq*) documents that *C5aR1* mRNA in normal mouse brain is predominantly expressed in microglia and macrophages. I. Flow cytometric plot shows that, after 3 days of AQP4-IgG infusion, greater numbers of neutrophils (Ly6G^+^ cells) infiltrate the lumbar parenchyma in WT mice than in *C5aR1* deficient mice. Percentages are quantified in the bar graph (*n* = 4 mice in each group). J. Quantification of the percentage of infiltrating neutrophils (CD45^+^CD11b^+^Ly6G^+^Ly6C^-^ cells among CD45^+^CD11b^+^ cells) in spinal cords of wild-type and *C5aR1^−/−^* mice at day 3 of AQP4-IgG infusion. K. Motor function, assessed by Rotarod performance (latency to fall), in WT and *C5aR1^−/−^* mice infused with AQP4-IgG or (only *C5aR1^−/−^* mice) normal mouse IgG (0.1 µg/µL, *n* = 6 mice per group). L. Motor function of mice assessed as fall latency from Rotarod; (time: *F* _(8, 48)_ = 1.543, *p*=0.1675; treatment: *F* _(1, 6)_ = 14.90, *p* = 0.0084; interaction: *F* _(8, 39)_ = 2.070, *p*=0.0629; *n* = 6-7 mice per group). In all graphs, data represent means ± SEM and all statistical tests are two-sided. One-way ANOVA, followed by Tukey’s *post hoc* multiple comparisons test in D. Unpaired Student *t*-test in J. Two-way repeated measures ANOVA with Sidak *post hoc* test in K. *p* < 0.05 was considered a significant difference.

Spinal gray matter is inflamed in patients with early stage NMO ^27–30^. To investigate the functional outcome for microglia activated by C5aR1 signaling in this mouse model, we evaluated microglial lysosomal activity in the lumbar ventral horn at the peak of neutrophil infiltration (day 3 of AQP4-IgG infusion). Application of mask and 3D Imaris rendering at the single cell level revealed prominent CD68^+^ phagolysosome structures within expanded microglial cytoplasm (Figure 5A; top panels). By contrast, the CD68^+^ lysosome was not expanded in *C5aR1*-null mice (Figure 5A; lower panels). The morphological alterations in spinal microglia on day 3 (quantitated in Figure 5B) presumably reflect their response to cues promoting lesion progression ^31^. Analysis by Imaris 10 AI-powered filament tracer revealed morphological alterations in the early response of WT mice to subarachnoid AQP4-IgG infusion, not seen in *C5aR1*-deficient recipients of AQP4-IgG, namely shorter microglial processes with less branch complexity (Figure 5C). The reduced complexity of processes in ventral gray matter microglia in the early stage NMO lesion coincided with phagolysosomal expansion. Neither change was observed in microglia of AQP4-IgG-infused *C5aR1*-knockout mice. Together, the results reveal that the C5a-C5a receptor 1 signaling pathway is a critical contributor to microglial activation in this model of NMO.

**Figure 5.**
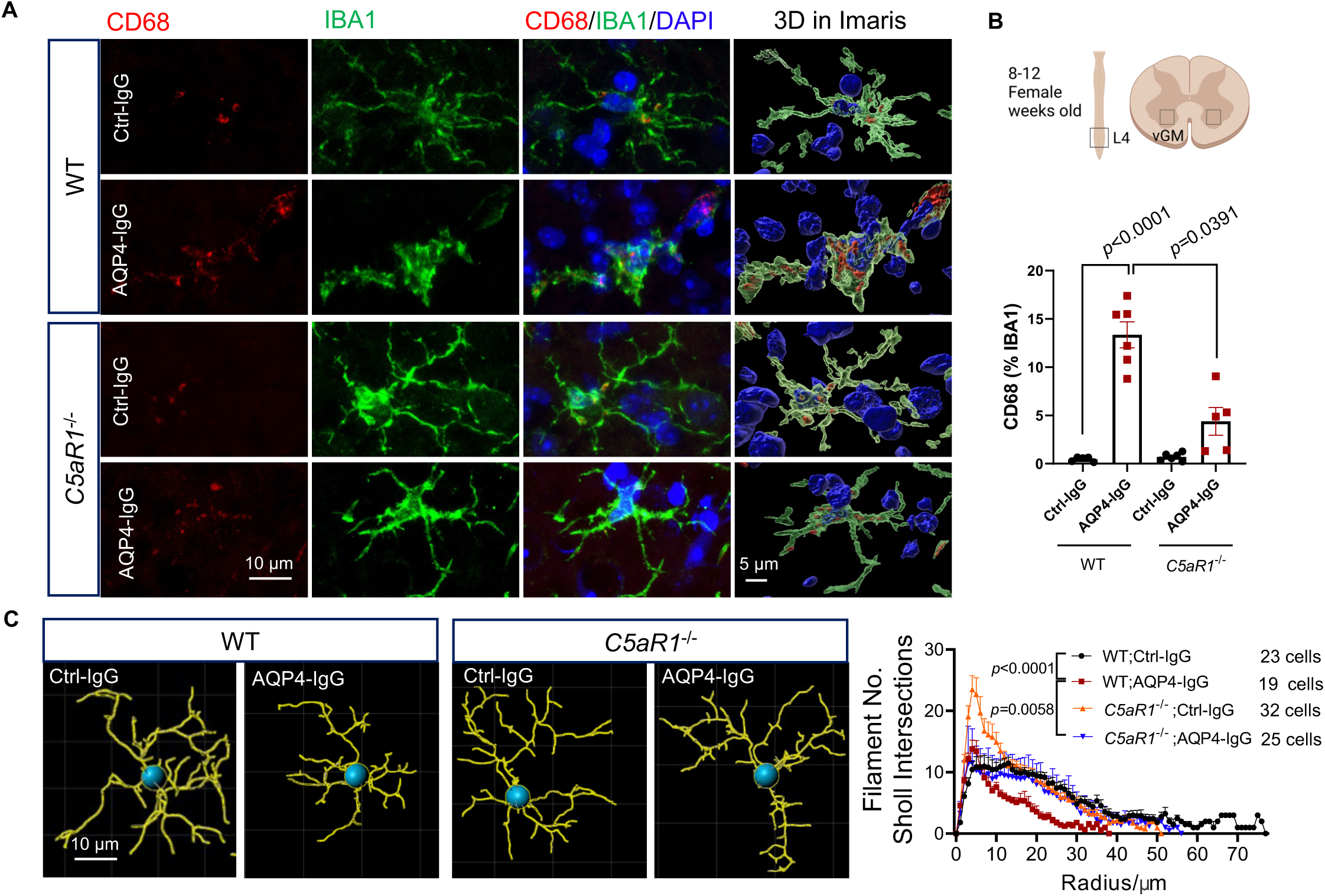
*C5aR1* deficiency abrogates downstream microglial activation response to AQP4-IgG infusion. A. Representative confocal images from lumbar cord of WT and *C5aR1*-deficient mice infused with normal IgG or AQP4-IgG. Lysosomal CD68-immunoreactivity (red) is more abundant in microglia (green) of WT recipients of AQP4-IgG than in *C5aR1*^−/−^ recipients. Imaris 3D rendering images illustrate the magnitude of lysosomal expansion. B. Image J analysis of the percentage area occupied by lysosome inside microglia in the lumbar cord of different experimental groups (Treatment: *F* _(1, 20)_ = 145.8, *p* < 0.0001; genotype: *F* _(1, 20)_ = 1.144, *p* = 0.2975; *n* = 5-6 mice per group). C. Sholl analysis of microglial branching revealed by Imaris AI-powered filament tracing which counts the number of microglial filaments intersected by 1 µm spherical steps (Treatment: *F* _(77, 3730)_ = 39.35, *p* < 0.0001; Radius: *F* _(3, 95)_ = 17.02, *p* < 0.0001; *n* = 19-32 microglia from 5 mice per group). In all graphs, data represent means ± SEM and all statistical tests are two-sided. Two-way (treatment × genotyping) ANOVA with Sidak *post hoc* multiple comparisons test in B. Two-way repeated measures ANOVA with Sidak *post hoc* test in C. *p* < 0.05 was considered significant difference.

### Lack of C5aR1 alleviates motor neuronal oxidative stress and lipid droplet accumulation

Functional impairment of spinal motor neurons causes weakness or paralysis in NMO attacks ^32^ and in rodent NMO models induced by intrathecal AQP4-IgG infusion ^16,33,34^. To better understand the pathophysiologic events linking motor neuronal dysfunction to initial astrocyte activation and secondary activation of neutrophils and microglia, we evaluated cytoplasmic Nissl body staining and choline acetyltransferase (ChAT) immunoreactivity and nuclear-cytoplasmic HuD immunoreactivity as indices of neuronal health during AQP4-IgG infusion. Loss of Nissl body and ChAT staining was observed in motor neurons on day 3 but, consistent with the behavioral data (Figure 3I and Figure S7), losses were transient and normalized by day 28 after stopping IgG infusion on day 7 (Figure 6A-D).

**Figure 6.**
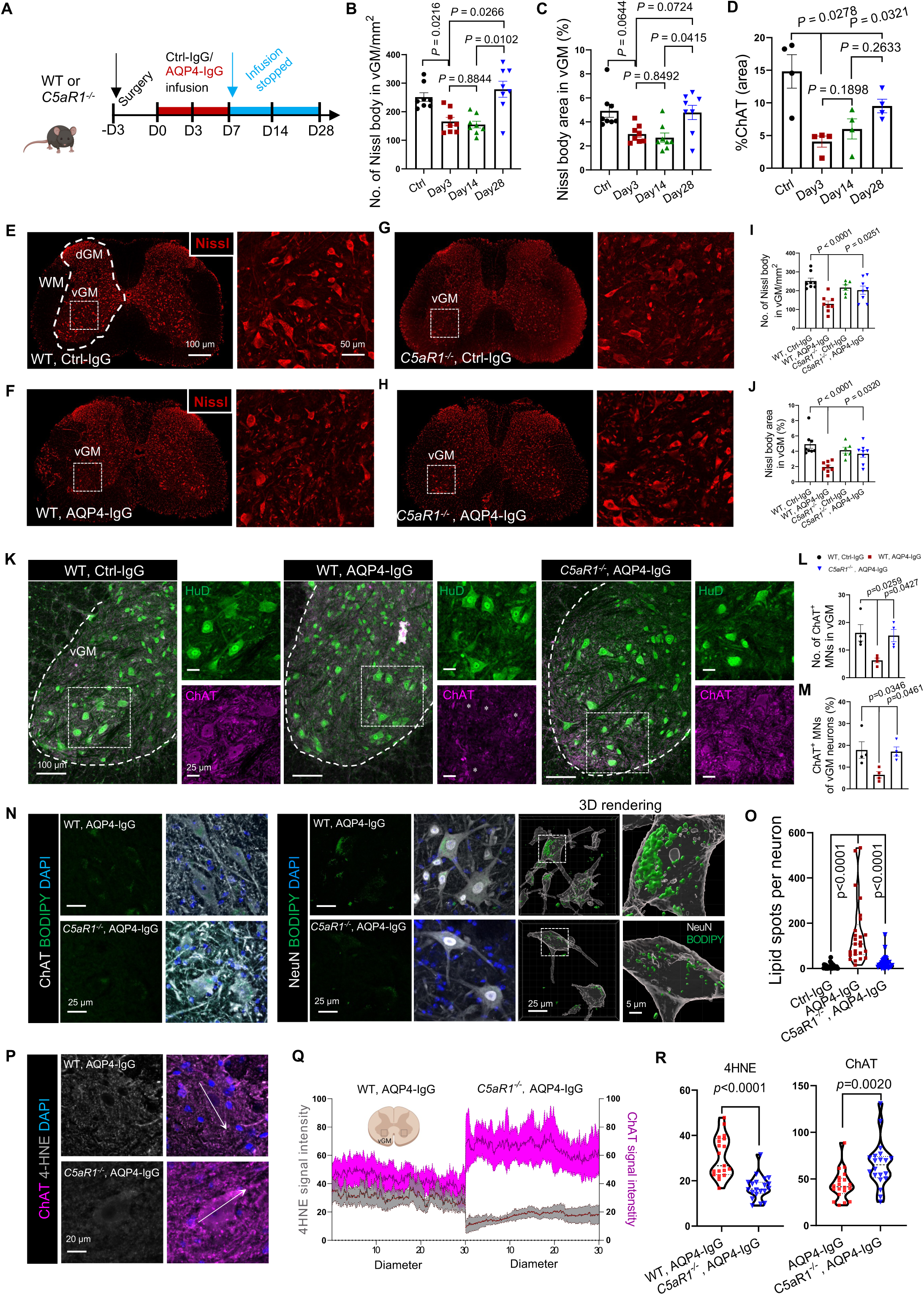
*C5aR1-*deficiency ameliorates neuronal dysfunction and lipid droplet accumulation in motor neurons. A. Experimental design: WT mice were infused continuously with AQP4-IgG using osmotic pumps from day 0 through day 7; infusion was stopped on day 8. B-D. Quantification of Nissl bodies (B), Nissl area (C) and percentage area occupied by ChAT^+^ motor neurons in grey matter of lumbar cord of Ctrl-IgG-infused mice at day 3, AQP4-IgG-infused mice at days 3, 14 and 28. Ctrl: (Nissl body count 251.5± 85.7/mm^2^, Ctrl Nissl body area: 4.928± 0.6 mm^2^, Ctrl ChAT signal area: 14.83 ± 1.8 mm^2^); Day 3: (Nissl body count 165.8 ± 21.6/mm^2^, Nissl body area 3.0 ± 0.4 mm^2^, ChAT signal area: 4.1 ± 0.8 mm^2^); Day 28: (Nissl body count: 278.8 ± 33.08/mm^2^, Nissl body area: 4.786 ± 0.6 mm^2^, ChAT signal area: 9.517 ± 1.8 mm^2^). E-H. Representative immunostained images of Nissl bodies (red) in ventral gray matter (vGM) neurons of wild-type and *C5aR1^−/−^* mice after 3 days’ Ctrl-IgG or AQP4-IgG infusion. Boxed areas are enlarged on the right. I, J. Numbers and sizes of Nissl body-positive neurons in ventral gray matter (*n* = 8 mice per group). K. Representative motor neuron confocal images and quantification in ventral gray matter of WT mice infused with Ctrl-IgG or AQP4-IgG, and *C5aR1^−/−^* mice infused with AQP4-IgG or control IgG (not shown). ANNA1[HuD]^+^ (green); ChAT^+^ (magenta). L, M. Numbers of ChAT^+^ motor neurons (MNs) and their percentage among total neurons (ANNA1 [HuD]^+^ in vGM. (*n* = 4 mice per group). N. Representative images of BODIPY^+^ lipid droplets (green) in ChAT^+^ motor neurons (grey) and NeuN^+^ neurons (grey) in ventral gray matter of AQP4-IgG-infused WT and *C5aR1^−/−^* mice. 3D reconstruction of BODIPY^+^ lipid and NeuN^+^ neurons using Imaris (right). O. Quantification of lipid droplet numbers in the cytoplasm of NeuN^+^ neurons in ventral gray matter of WT and *C5aR1* deficient mice infused for 3 days with AQP4-IgG (*n* = 20 neurons from 4 individual mice per group). P. Representative confocal images of 4HNE (peroxidative stress marker) in ChAT^+^ motor neurons of WT and *C5aR1^−/−^* mice infused with AQP4-IgG. Q and R. Q, Quantification of 4HNE and ChAT immunoreactivity intensities across 30 μm neuronal diameter; R, presented as mean ±SEM (*n*=20 neurons from 4 individual mice per group). All data represent means ± SEM, all statistical tests are two-sided. Two-way (treatment × genotyping) ANOVA with Sidak *post hoc* test multiple comparisons test in I and J. One-way ANOVA with Tukey *post hoc* in B-D, L and M. Unpaired Student *t*-test in O and R. *p* < 0.05 was considered significant difference.

On day 3, Nissl bodies in ventral gray matter neurons exhibited a 41% reduction in both number and average area (Figure 6 E, F, I and J); numbers of neurons staining for ChAT (exclusive to motor neurons) were 75% reduced and the percentage of ChAT^+^/HuD^+^ neurons was 70% reduced in AQP4-IgG-infused mice (Figure 6K-M). However, total numbers of HuD^+^ neurons in ventral gray matter did not differ significantly in control-IgG and AQP4-IgG-recipient mice (Figure S8 A-D). AQP4-IgG recipient mice lacking C5aR1 did not exhibit Nissl body loss (Figure 6I, J). These observations implicate neutrophil-microglial signaling through C5aR1 in the mediation of motor neuron dysfunction in this mouse NMO model. We next evaluated large ChAT-containing ventral horn neurons for morphologic evidence of dysfunction downstream of AQP4-IgG binding to astrocytes. The 75% reduction in numbers of motor neurons expressing ChAT suggests that the observed Nissl body loss reflects impaired motor neuron function, rather than actual neuronal loss. In mice with genetically-ablated C5aR1 signaling, motor neuronal ChAT immunoreactivity was retained at an apparently critical early stage of NMO evolution. The lack of significant difference in numbers of ventral horn interneurons (HuD^+^/ChAT^-^) between wild-type mice receiving control IgG or AQP4-IgG (Figure S8E) implies an inherent susceptibility of spinal motor neurons (HuD^+^/ChAT^+^) to functional impairment in NMO.

We investigated next whether disruption of astrocyte function, prior to terminal immune-mediated lysis, might impact neuronal lipid metabolism. Application of the neutral lipid stain BODIPY 493 revealed early accumulation of cytosolic lipid droplets in motor neurons during AQP4-IgG infusion (Figure 6N, O). Significantly fewer lipid droplets were found in corresponding neurons of control-IgG recipient mice (data not shown). To determine whether lipid droplet accumulation paralleled lipid peroxidation in motor neurons, we immunostained for the oxidative stress product, 4-hydroxynonenal (4-HNE). Its enhancement coincided with lipid droplet accumulation (*p* < 0.0001 for 4-HNE, *p =* 0.0020 for ChAT, Figure 6P, Q and R). Importantly, lipid peroxidation and droplet accumulation in spinal motor neurons did not differ significantly in *C5aR1**-***deficient recipients of AQP4-IgG and WT recipients of control IgG. Thus, neuronal lipid droplet accumulation and lipid oxidation in neurons is a C5aR1-signaling-dependent pathophysiological phenomenon occurring early in the evolving NMO lesion, independent of cytolytic complement lesioning.

### Microglia are required for AQP4-IgG to induce astrocytic production of CXCL1

Early experimental models of NMO, both *in vitro* ^26^ and *in vivo* ^35^, implicated a pathogenic role for granulocytic CNS infiltration. Astrocyte-derived CXCL1 is known to drive neutrophil transmigration in mouse models of viral encephalitis ^36^ and ischemic stroke ^37^. To investigate whether CXCL1 protein production is upregulated following astrocytic AQP4 ligation and internalization by IgG (Figure 7A, upper), we incubated WT or *AQP4-*null mouse astrocytes with pathogenic or non-pathogenic monoclonal AQP4-IgGs; IFN-γ plus TNF-α served as a control activator of astrocytes (Figure 7A, lower). *AQP4-*null mouse astrocytes were confirmed AQP4 deficient by immunoblotting (Figure 7B). Compared to the effect of non-pathogenic C-terminal cytoplasmic domain-reactive AQP4-IgG (CCD-m5, control), the immunofluorescence staining intensity of cytoplasmic CXCL1 was increased 244-fold (*p* = 0.0112, Figure 7C, D, upper) after 48 hrs’ exposure to the pathogenic AQP4-IgG (ECD-m21, extracellular domain-reactive) and secreted CXCL1 levels (detected by ELISA) rose significantly (1305 pg/mL *vs*. 159 pg/mL, *p* = 0.0030, Figure 7C, D, lower). However, in wild-type glial cultures depleted of microglia, pathogenic AQP4-IgG increased CXCL1 production only 8-fold (Figure S9) and the cytoplasmic CXCL1 increase was significantly attenuated in *AQP4*-null astrocytes co-cultured with microglia (Figure 7D; upper, *p* = 0.0164; lower, *p* = 0.0011). These findings indicate that upregulation of astrocytic production and secretion of CXCL1 by AQP4-IgG requires microglia-astrocyte interaction.

**Figure 7.**
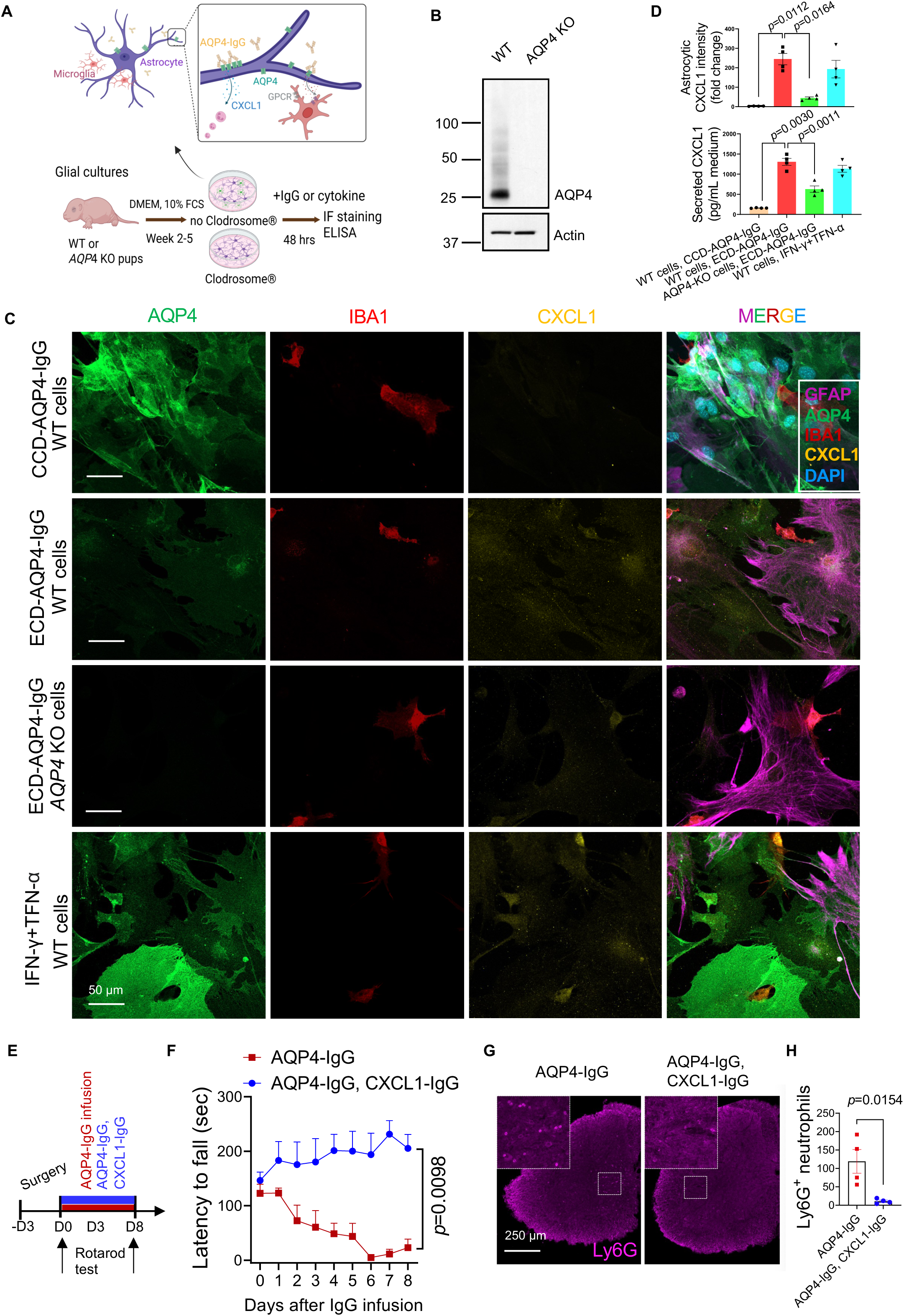
Microglia are required for pathogenic AQP4-IgG to upregulate CXCL1 in cultured mouse astrocytes. A. Experimental design and schematic representation of the sequelae of IgG binding to AQP4 on astrocytes in primary glial cultures established from wild-type and *AQP4*-null pup brains. Microglia were depleted by treating Clodrosome (100 μg/mL) before subculture. Two days after adding IgG or cytokines, CXCL1 production was assessed by immunostaining or ELISA. B. SDS-PAGE and immunoblot analysis of WT, *AQP4*-KO primary glial cells using antibodies to mouse AQP4, and actin (loading control). C. CXCL1 immunoreactivity was assessed in wild-type and *AQP4* knockout glial cells exposed to a control non-pathogenic monoclonal mouse IgG specific for the AQP4 cytoplasmic C-terminal domain (CCD-AQP4-IgG, m5), a pathogenic extracellular domain-reactive IgG (ECD-AQP4-IgG, m21) or to IFN-γ plus TFN-α cytokines. Astrocytes are identified by AQP4 and GFAP immunoreactivities; microglia by IBA1; DNA is blue (DAPI). D. Cellular CXCL1 (upper) was quantified from fluorescence intensity in C images; secreted CXCL1 protein levels (lower) were quantified in the glial culture media by ELISA. E. Experimental design: two groups of WT mice were infused continuously via spinal subarachnoid catheter from osmotic pump (day 0 to day 8) with AQP4-IgG or AQP4-IgG mixed with CXCL1-IgG. F. Motor function of mice in E was assessed as fall latency from Rotarod; (time: *F* _(8, 40)_ = 1.319, *p* = 0.2632; treatment: *F* _(1, 5)_ = 16.38, *p* = 0.0098; interaction: *F* _(8, 22)_ = 7.766, *p* < 0.0001; *n* = 4-5 mice per group). G-H. Representative confocal images at day 7 of IgG infusion and quantification of neutrophils (Ly6G^+^) in lumbar cord parenchyma (*n*= 4 mice per group). Higher magnifications (left corners) of boxed areas in G show neutrophils (magenta). In all graphs, data represent means ± SEM; all statistical tests are two-sided. One-way ANOVA with Tukey *post hoc* in D; *n* = 4 wells in D. Two-way repeated measures ANOVA with Sidak *post hoc* test in F. Unpaired Student *t*-test in H, *p* < 0.05 was considered a significant difference.

To investigate whether blockade of CXCL1-mediated granulocytic trafficking *in vivo* might attenuate motor dysfunction in this mouse NMO model, we co-infused anti-CXCL1-IgG together with AQP4-IgG or control normal mouse IgG into the subarachnoid space for 7 days (Figure 7E). Behavioral testing revealed that inhibition of CXCL1 signaling prevented motor impairment by AQP4-IgG infusion. The motor performance of mice treated with anti-CXCL1-IgG (Figure 7F and Movie S 5) was comparable to that of neutrophil-depleted mice (Figure 3G). Thus, CXCL1-dependent neutrophil recruitment is a critical step in initiating an NMO attack.

### A novel (Galectin3/P2Y12-positive) subpopulation of disease-associated microglia interacts with motor neurons

To investigate how microglia-neuron interactions might modulate neuronal function, we analyzed the surface phenotype of microglia in the ventral horn of AQP4-IgG–infused mice by immunostaining microglial IBA1 and motor neuronal ChAT. Two-dimensional and three-dimensional analyses revealed a significant increase in microglial contacts with somata of motor neurons (Figure 8A-C) by comparison with control-IgG infused mice. Galectin-3, a marker of disease-associated microglia (DAMs), was highly expressed on microglia engaged with neurons. The Galectin-3 signal area was 122% greater in microglia that were contacting neurons compared to microglia not contacting neurons (Figure 8D). Microglia are known to regulate neuronal activity through sensing neuron-derived ATP/ADP via the P2Y12 purinergic receptors enriched on their processes ^31,38–40^. In the context of “NMO” disease evolution in AQP4-IgG– recipient mice, we found that 38.5% of the Galectin-3^+^ microglial clusters interacting with motor neurons highly co-expressed P2Y12 receptors. In comparing immunohistochemical changes in spinal cord tissue of NMO patients, microglia associated with motor neurons were found to be P2Y12^+^ (Figure 8H). Together, these findings identify a previously unrecognized microglial subpopulation expressing both a DAM marker (Galectin-3) and a homeostatic marker (P2Y12).

**Figure 8.**
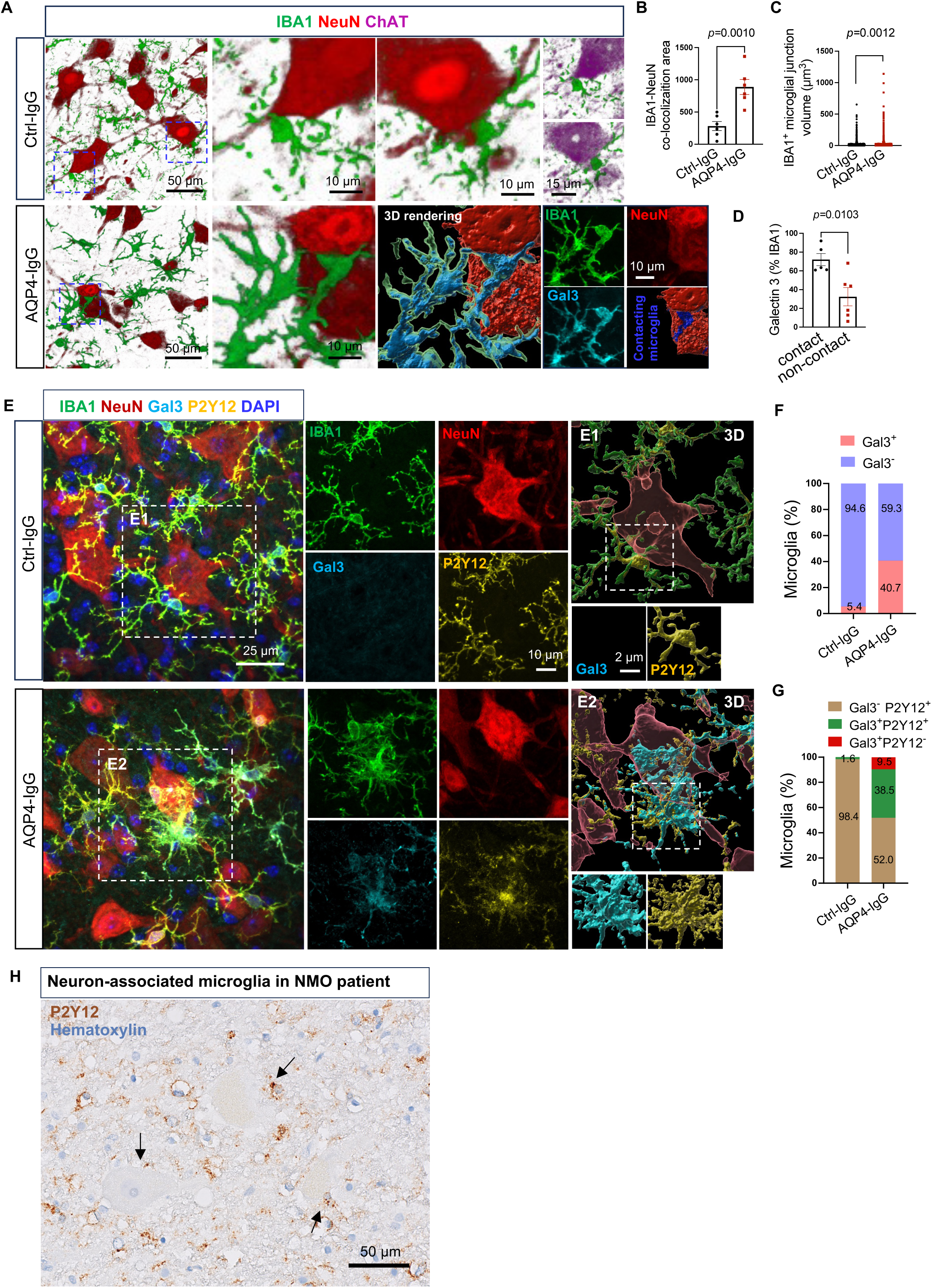
Disease-associated microglia (DAM) interact with motor neurons in NMO mouse model. A. Representative immunostained images of motor neuron (ChAT^+^)-associated microglia (IBA1^+^) in ventral gray matter of mice after 3 days’ Ctrl-IgG or AQP4-IgG infusion. Boxed areas are enlarged on the right. 3D rendering of boxed area in AQP4-IgG mice is shown as split confocal channels (lower right) and merged (lower center). B and C. Quantification of microglia-neuron co-colocalization area (B) and volume (C) in A (*n* = 6 mice per group in B; *n* = 1526 and 1272 microglial contacts, respectively, for Ctrl-IgG and AQP4-IgG mice). D. Image J analysis shows the percentage area occupied by Galectin 3 within contacting microglia is greater than in non-contacting microglia in lumbar cord of the two experimental groups. E. Overlaid confocal images show microglial P2Y12 receptor (green), Gal3 marker of DAMs (Cyan) contacting neurons in lumbar spinal ventral gray matter of AQP4-IgG-infused mice, in contrast to control-IgG-infused mice. F. Percentage of DAMs (IBA1^+^Gal3^+^) contacting neurons is significantly increased after AQP4-IgG infusion. G. The enhanced expression of P2Y12 receptor by IBA1^+^Gal3^+^ DAMs (38.5%) contacting neurons implicates P2Y12 in the contact mechanism. H. Immunohistochemistry reveals neurons associated with microglia (P2Y12^+^, brown) in this early-stage lesion in the spinal cord ventral gray matter of an NMO patient. Arrows identify microglia-neuron contact sites.

## DISCUSSION

Neutrophil infiltration is a histopathological feature that distinguishes NMO from other inflammatory CNS demyelinating disorders ^5^. Our findings in a mouse model of NMO assign neutrophils a critical, therapeutically targetable, role in initiating motor neuronal dysfunction prior to activation of the classical complement cascade and ensuing irreversible tissue destruction. Analogous to observations in early CNS lesions of NMO patients ^8^, spinal subarachnoid infusion of a pathogenic, non-complement-activating monoclonal AQP4-IgG into mice causes neutrophil migration into the cord parenchyma to undergo NETosis. Our initially reported mouse NMO model identified a C3a-C3aR1 signaling-dependent interaction between AQP4-IgG-activated astrocytes and microglia as a driver of initial lesion formation and motor impairment ^16^. Our present study demonstrates that neutrophils are recruited soon after *in vivo* engagement of pathogenic IgG with astrocytic AQP4, with neutrophil-microglia interaction via C5a-C5a receptor signaling being a critical co-stimulatory event for the development of motor paresis. Surprisingly, we found no C5 protein in the evolving CNS lesion, but the cytoplasm of infiltrating neutrophils contained abundant C5a immunoreactivity. We concluded that C5 was synthesized by activated neutrophils, rapidly cleaved post-translationally by cytoplasmic cathepsins and secreted as C5a ^41,42^. Thus, our study suggests the complosome of CNS-infiltrating neutrophils is a non-canonical source of C5a synergizing with astrocyte-derived C3a^16^ to co-stimulate microglia and serving as a cell-autonomous mechanism enhancing neutrophil influx.

In the context of aging and neurodegenerative diseases, neuronal oxidative stress induces toxic lipid droplets accumulation in astrocytes ^43^. This is attributed to impaired transfer of lipid products from neuron to astrocyte for mitochondrial catabolism by β-oxidation ^44^. Immunohistopathological studies of sublytic astrocytopathy in CNS tissues of NMO patients suggest that damage signals from lesional astrocytes are propagated via the panglial syncytium to distant astrocytes that are not themselves ligated by IgG (i.e., they retain surface AQP4) ^45^. It is conceivable that microglia, fully activated by astrocytic C3a ^16^ and neutrophilic C5a (present data), are primed to release factors that further promote astrocytopathy, such as by impairing the processing of lipids transferred from neurons. Important observations in our present study are 1) neurochemical evidence of motor neuronal oxidative stress, loss of choline acetyltransferase and lipid droplet accumulation accompanying initial clinical motor impairment that is reversible on removal of pathogenic AQP4-IgG, and 2) motor dysfunction preceding endothelial tight junction disruption. We therefore propose that the hallmark complement-mediated cytodestructive end-stage lesions responsible for irreversible NMO disability require gross blood-brain barrier disruption allowing astrocytic membrane lesioning by deposition of plasma-derived opsonic macromolecules, including IgM and C1q ^5^. It is at this point that neutralization of circulating and CNS-penetrating C5 proenzyme by anti-C5 monoclonal antibody therapy (eculizumab) is highly efficacious in aborting an acute attack and preventing NMO relapse ^46^. Our study demonstrates that C5a/C5aR1 signaling between infiltrating neutrophils and resident microglia (the predominant C5aR1-expressing cells in the CNS) is an early pathogenic mechanism contributing to neuronal stress and motor dysfunction.

We discovered several cellular mechanisms that plausibly underlie both the evolving histopathological lesion and motor impairment. Neutrophil-attractant chemokines (CXCL1, CXCL5, CXCL7, CXCL8) and activation markers (MPO and NE) are elevated in attack phase CSF of NMO patients ^47,48^. Our glial culture experiment demonstrated that AQP4-IgG stimulates astrocytic production and release of CXCL1. The far lower CXCL1 levels induced by IgG in *AQP4*-null astrocytes and in microglia-depleted cultures indicate this mechanism is both antigen-dependent and microglia-dependent. It is yet to be determined whether CNS infiltration by neutrophils is solely dependent on CXCL1 secretion by activated astrocytes.

Recognition by clinicians that neutrophils and their chemokines aid the distinction of NMO from multiple sclerosis ^7^, and that neutrophil infiltration correlates with lesion size in NMO patients ^8^, underscores the therapeutic relevance of our findings. The demonstration by Saadoun et al ^9^ that administration of neutrophil protease inhibitors limited the lesion size in mice with disrupted blood-brain barrier (injected intracerebrally with NMO patient IgG combined with fresh human complement) highlighted the significance of neutrophils as therapeutic targets in the cytodestructive phase of NMO. Our study has revealed a critical earlier role for neutrophils in initiating NMO pathophysiology prior to tissue damage.

Targeting of neutrophils or C5a receptors is an attractive therapeutic option for attack intervention in NMO. Avacopan, a C5aR1 antagonist, is FDA-approved for treating anti-neutrophil cytoplasmic antibody-associated vasculitis ^49^. The blood-brain barrier-permeable C5aR1 inhibitor, PMX 53, was demonstrated to prevent astrocytic lysis in rats co-injected intracerebrally with AQP4-IgG and human serum as an exogenous source of complement components ^35^. By demonstrating that motor paresis can be mitigated by the C5aR1 antagonist, PMX 205 in the precytolytic phase of NMO, our study highlights the critical pathophysiological importance of neutrophil-microglia interaction preceding activation of the classical complement cascade and resultant cytodestruction.

## LIMITATION

Release of proteolytic enzymes, bioactive lipids, and reactive oxygen species by degranulating neutrophils is considered to cause neuronal oxidative stress, and may indeed contribute to motor neuron dysfunction in NMO ^50–52^, but those events occur later than the time frame of neurological impairment that our study has addressed. Activated astrocytes ^36^ and secondarily activated neural, endothelial, T cells and myeloid cells, both resident and infiltrating, all undoubtedly contribute to NMO pathology ^53^. However, the molecular contributions of microglia to oxidative stress and lipid droplet accumulation in motor neurons remain poorly understood, as does the basis for selective vulnerability of motor neurons. Our preliminary identification of a novel Galectin3^+^/P2Y12^+^ subpopulation of disease-associated microglia contacting motor neurons provides a new direction for future investigations ^54^. Spatial and single-cell proteomics should further elucidate the complex neutrophil–microglial interactions driving early NMO pathogenesis.

## MATERIALS AND METHODS

### Mice

We used 8-12-week-old female C57BL/6J wild-type, *Cx3cr1^GFP/^*^+^ transgenic mice, *C5aR1^−/−^* mice. C57BL/6J (stock no. 000664), *Cx3cr1^GFP^*^/*GFP*^ (B6.129P2(Cg)-*Cx3cr1^tm1Litt^*/J, stock no. 005582), *C5aR1^−/−^* (B6.129S4(C)-*C5ar1^t^*^m1Cge^/BaoluJ, stock no. 033903), *CCR2^CreER-GFP^*(C57BL/6-*Ccr2*^em^^1(icre/ERT^^2)Peng^/J, stock no. 035229) mouse lines were purchased from The Jackson Laboratory and bred at Mayo Clinic. Mice were group-housed on a 12-hour light/dark cycle and a temperature– and humidity-controlled room. Animal procedures and protocols were approved by the Mayo Clinic Institutional Animal Care and Use Committee (IACUC protocol no. 2909-17) and were conducted in agreement with the National Institutes of Health (NIH) Guide for the Care and Use of Laboratory Animals. *Cx3cr1^GFP/^*^+^ mice were used to visualize microglia and neutrophil interaction for electron microscopy imaging.

### Spinal catheter placement and IgG delivery

Under isoflurane anesthesia, we inserted into the subarachnoid space via the cisterna magna (through a 1 mm horizontal incision in the dura mater) a 6-cm polyurethane intrathecal catheter (ALZET, no.: 0007743), facilitated by a Teflon-coated stainless-steel stylet, and extended it to the lumbar spinal level. Dental glue was applied to prevent leaks from catheter connections, and the catheter was anchored by suturing to neck musculature. On awakening, mice were observed postoperatively for 2 hrs for motor signs of traumatic cord injury. Injured mice (∼10%) were promptly euthanized. After 3 days’ training on the Rotarod beam (days –3 to –1, 4 rpm/min, 5 min) with daily slow manual catheter flushing with artificial CSF, an osmotic minipump delivery system (ALZET, 1007D), containing a pathogenic mouse monoclonal anti-AQP4-IgG1 (mECD, AQP4-m21, created in-house; licensed to Sigma as MABN2471) or mouse polyclonal serum IgG (Sigma, I5381), or anti-AQP4-IgG plus rat monoclonal anti-CXCL1-IgG (R & D, MAB453, 0.5 μg/10 μL/day for 8 days) was placed subcutaneously between the shoulders. AQP4-IgG or mouse serum IgG was infused continuously for 1, 3 or 7 days (1.2 μg/12 μL/day).

### Rotarod training and testing

Daily testing began day 0, at 4 rpm/min and accelerated to 40 rpm/min over a 5-minute period. Each mouse underwent 3 trials at 15-minute intervals. The recorded latency to fall from the Rotarod wheel was averaged for each mouse.

### Tissue collection and immunofluorescence staining

Spinal cord tissues were harvested following euthanasia and transcardiac perfusion with 30 mL ice-cold PBS. After 10% formalin perfusion, fixation (16-24 hrs at 4), then cryoprotection by immersion in 30% sucrose), tissues were cryosectioned coronally (16-40 µm thick, Leica cryostat CM1520) and embedded in Tissue-Tek OCT compound (Figure 1 and 4); 50 µm thick sections were prepared by vibratome (LEICA VT1000S) without embedding (all other Figures) for immunofluorescence staining. We mounted 16 µm sections directly onto SuperFrost Plus microscope slides (stored at –80) and prepared 40 µm sections for free-floating staining (stored in 30% glycol). For immunofluorescence staining, sections were permeabilized and blocked (30 min), using 0.25% Triton X-100 in PBS containing 2.5% bovine serum albumin and 5% donkey serum, then exposed 16-24 hrs at 4 to the following primary antibodies in PBS containing 0.1% Triton X-100 (Sigma) and 1% BSA: rat anti-Ly6G (Bio X Cell; BE0075-1, 1:500), Human/Mouse goat anti-myeloperoxidase/MPO (R & D; AF3667-SP; 1:800), rat anti-neutrophil elastase/NE (R & D, MAB4517-SP; 1:500), mouse anti-C5 (Hycult Biotech; HM1073, 1:500), rat anti-C5a (Invitrogen; MA5-23910, 1:300) and goat anti-GFP (Abcam; Ab6673, 1:1,000), goat anti-CD31 (R & D; AF3628, 1:500), rabbit anti-IBA1 (Abcam; Ab104225, 1:500), goat anti-IBA1 (Wako; 011-27991, 1:500), guinea pig anti-IBA1 (Synaptic Systems; 234 308, 1:1,000), rabbit anti-ChAT (Sigma-Aldrich, AB143; 1:100), rabbit anti-P2Y12 (AnaSpec; AS-55043A; 1:1,000), human anti-HuD ^55^, from patients with paraneoplastic neurological autoimmunity related to small-cell lung carcinoma; 1:10,000), goat anti-human/mouse/rat galectin-3 antibody (R & D; AF1197, # NP_034835, 1: 1,000) and mouse anti-4-Hydroxynonenal/4-HNE (R & D; MAB3249-SP, clone # 198960, 1:200). Sections were then exposed (22, 2 hrs) to appropriate fluorochrome-conjugated secondary antibodies (diluted in PBS containing 0.1% Triton X-100 and 1% BSA) and then counterstained with DAPI (Sigma-Aldrich, 0.1μg/mL) and mounted with SlowFade™ Diamond Mountant (Thermo Fisher Scientific). The slides were held at 22 overnight in the dark to set the mounting medium. Images were acquired using either a ×63 oil-immersion objective lens or a ×40 water-immersion objective lens with a Zeiss LSM980 confocal microscope, ensuring optimal resolution for different experimental setups. For electron microscopy (EM) imaging of neutrophil-microglia interaction, the cord was expelled from the vertebral canal post-PBS perfusion using ice-cold PBS from a 3-mL syringe connected to a 200 µL pipette. Lumbar cord (2 mm-length) was quickly immersed (2 hrs at 4) in 4% EM grade paraformaldehyde aqueous solution (Electron Microscopy Sciences, no. 15710, diluted 1:4 in PBS). Vibratome sections were prepared without embedding (50-100 µm, LEICA VT1000S). The immunostaining method was standard except that: 1) primary antibodies (rat anti-Ly6G, 1:200, and rabbit anti-NeuN, 1:500), and Alexa fluor-conjugated secondary antibodies (donkey anti-rat IgG, 594, and donkey anti-rabbit IgG, 405) were diluted 1:500) in PBS without permeabilization; 2) secondary antibody exposure was at 4°C for 24 hrs, with mild shaking; 3) cord sections were mounted in 30% glycerol on glass slides for confocal imaging. For EM imaging of lipid-droplets, lumbar cord (2 mm-length) was quickly dissected and postfixed in 2% glutaraldehyde and 2% paraformaldehyde in 0.1 M cacodylate buffer containing 2 mM calcium chloride without immunostaining. Surface rendering, spot rendering, filament rendering and visualization were performed using Imaris v.10.0 (Oxford Instruments).

### Immunohistochemical analysis of human spinal cord

An archival block of spinal cord from an autopsied NMO patient was subjected to neuropathological evaluation, as previously detailed ^45^. Sections (5 μm) of paraffin-embedded formalin-fixed tissue were stained with hematoxylin-eosin, Luxol fast blue-Periodic acid Schiff (LFB/PAS) myelin stain. Immunohistochemistry utilized an EnVisionTM FLEX + secondary system (Dako/Agilent) ^56^. Primary IgG antibodies included rabbit anti-AQP4 (Sigma; A5971; 1:250), KiM1P (monoclonal, gift from Dr. W. Bruck, Germany), and rabbit anti-human P2Y12 (Abcam; ab300140; 1:1,000). Images (×40) were captured using an Olympus BX51 microscope (EVI-DENT, Tokyo).

### Single-cell suspension for flow cytometry

Spinal cords from three cohorts of mice were perfused and harvested after 1-, 3-, and 5-days’ subarachnoid infusion with AQP4-IgG; cord tissue from a control cohort infused with normal mouse serum IgG was harvested on day 3. All mice were terminally perfused with ice-cold PBS. Lumbar cords were rapidly harvested, digested in HBSS containing collagenase I/IV and DNase and shaken (New Brunswick Scientific NB-C24; 37, 300 rpm, 35 min), with trituration by polished glass pipette every 10 min to aid dissociation ^57^. After passage through 70 µm nylon mesh (no. 130-110-916), the suspension was centrifuged (515*g*, 7 min at 4□), and the pellet was resuspended in PBS, gently overlaid on 30% Percoll and centrifuged (800*g,* 20 min at 4□). Sedimented cells, free of debris and myelin, were resuspended in ice-cold PBS, counted, filtered via 70 μM cell strainer (Falcon no. 352052), immunostained and phenotyped by flow cytometry.

### High-parameter flow cytometry and analysis

A preliminary experiment determined optimal antibody dilutions and appropriately set compensation for the antibody panel’s multiple fluorochromes ^58^. After resuspension in PBS, cells were first stained with Zombie UV viability dyes (BioLegend, 1:1,000; 20 min at room temperature in the dark). Fc-receptors were blocked using rat IgG2b anti-mouse CD16/CD32 (BioLegend, 101302, 1:50) and the following mixed panel of surface-staining antibodies was applied (30 min, 4□): CD45-PE-CF594 (BD, 562420, 1:1,000), CD11b-PE-Cy5 (Tonbo, 55-0112-U100, 1:1,000), CD11c (BioLegend, 1:200), CD4-BV510 (BioLegend, 100449, 1:200), CD8a-BV785(BioLgend, 100750, 1:200), CX3CR1-Pacific Blue (BioLegend, 149038, 1:300), CD206 (1:200), Ly6G, Ly6C-PerCP (BioLegend, 128028, 1:100□), CD3-PE-Fire700, TMEM119, C5aR-PE (BioLegend, 135805, 1:100), NK1.1-APC/FireTM 810 (BioLegend, 156519, 1:200), MHC-II (BD, 746197, 1:300), P2ry12-APC (BioLgend, 848005, 1:500), B220-Spark NIR685 (BioLgend, 103268, 1:200). After washing, the samples’ emission spectra were measured on a 5-laser Aurora cytometer (Cytek; Spectro Flo software). The data were analyzed using FlowJo™ v10.10 Software (BD Life Sciences).

### Near-infrared branding (NIRB)

We acquired whole mount confocal images of sections with ×20 lens, tiles and Z-stack, then enlarged (×63 oil lens with Z-stack) foci of interest containing neutrophils adjacent to microglia (Ly6G/Cx3cr1-GFP). The spinal cord section was then rinsed and plated in 1×PBS on a new glass microscope slide. We generated fiducial branding marks on spinal cord sections by progressively increasing the laser power of a two-photon microscope at micrometer-precision points. The holes created by induced microbubbles enabled later reidentification of the field containing microglia and neutrophils of interest. The section was next postfixed (4, 24 hrs) in 2% glutaraldehyde and 2% paraformaldehyde in 0.1 M cacodylate buffer containing 2 mM calcium chloride.

### Serial block-face scanning electron microscopy (SBF-SEM)

SBF-SEM was performed as previously described ^59^. We used a workflow for combining the two techniques to allow detection of individual regions of interest (ROIs) in a large field at small X, Y pixel size, and then imaged the subsequent targeted volume at high voxel resolution. EM imaging was acquired under high vacuum/low water conditions with a starting energy of 1.8 keV and beam current of 0.10 nA. Stacks of several hundred sections, 100 nm thick, were obtained by the diamond knife and imaged at 15 nm × 15 nm × 100 nm spatial resolution. In alignment with Z-stack confocal images, the targeted microglia-neutrophil couple can be reconstructed based on the automatically obtaining EM-level 3D database. For analysis, Fiji-Image J (NIH, Bethesda, MD) and Reconstruct software packages were used to identify cell morphology, interaction, ultrastructure and cell segmentation.

### 3D image segmentation

Image segmentation, volume rendering and visualization were conducted using Reconstruct software packages ^60^. Neuron, microglia, and neutrophil morphology were identified in every 100-nm-thick section based on confocal imaging. For neutrophil segmentation, multi-lobed nuclear structure and types of membrane-bound cytoplasmic granules were the main neutrophil identifying characteristics ^61^. The neutrophil boundary was outlined based on darker cytoplasm compared to that of microglia. For microglial segmentation, we observed a relatively dark nucleus with clumped chromatin and lysosomes in lighter cytoplasm ^62^. Neurons were defined by clear nucleus with prominent nucleoli and a distinctive soma with abundant endoplasmic reticulum ^63^.

### Sholl analysis using Imaris filament tracer

Microglial branch analysis of IBA1-stained lumbar anterior horn tissue was conducted using confocal Z-stack images acquired with a × 63 oil immersion lens (Zoom × 0.6, 2,048 × 2,048 pixels, 0.082 μm per pixel, 0.5 μm Z-step, 40 slices). Imaris software (v.10.0, Oxford Instruments) was utilized to render and quantify 3D microglial branches via the “Filament tracing” module (https://imaris.oxinst.com/versions/10). For analysis, we examined 3-5 randomly selected cropped 3D volumes per confocal image. The microglial soma diameter was set to 6 μm based on DAPI staining, and the minimum process detail was determined to be 0.35 μm. The tracing workflow included soma detection, seed point identification along cellular processes, and computation of filamentous segments. Segments were classified as “good” or “bad” by comparison with confocal images. To enhance efficiency and accuracy, we employed an AI-powered filament tracer integrated with an intensity-based segmentation approach to differentiate genuine microglial processes from background. A training set of ∼100 manually annotated seed points per group were generated for machine learning-based classification, which was iteratively refined through an automated classification process. For process intersection quantification, we generated concentric spheres centered on the soma, increasing in 1 μm increments from a 6 μm radius. Intersection values at each radius were calculated using “Filament No. of Sholl intersections” model in Imaris.

### CD68 imaging and analysis

Lumbar cord sections were incubated overnight at 4 with rabbit-CD68 (Abcam, ab125212, 1:500) and goat anti-IBA1 (Wako; 011-27991, 1:500), diluted in PBS containing 0.25% Triton X-100. Following primary antibody incubation, sections were washed three times in PBS (5 min each) and stained 2 hrs with Alexa 594 donkey anti-rabbit (1:1,000) and Alexa 488 donkey anti-goat IgG (1:1,000). Nuclei were counterstained with DAPI (Sigma, 0.1 μg/mL) and mounted with SlowFade™ Diamond Antifade Mountants (Thermo Fisher Scientific). The CD68 expression in the ventral grey matter was analyzed using a LSM980 confocal microscope with ×40 water objective lens (20 z slice, 0.5 μm per slice). CD68 fluorescence (excitation/emission: 561 nm/590-618 nm) and IBA1 fluorescence (488 nm/496-525 nm) were captured. Binary CD68 images were generated using a uniform threshold in ImageJ, and CD68 expression was quantified from maximum-intensity Z-projections. The percentage of CD68 expression was calculated as the CD68^+^ area relative to the IBA1^+^ area. 3D renderings of CD68^+^ puncta, IBA1^+^ microglia, and DAPI^+^ nuclei were generated using the “Surface” model function in Imaris v10.0, with transparency adjustments applied to microglial surface rendering for enhanced visualization and spatial analysis.

### Pharmacological inhibition of C5aR1 signaling in *vivo*

Trifluoroacetate was removed from the cyclic hexapeptide C5aR1 antagonist PMX 205 (HY-110136 A; MedChemExpress, NJ, USA) by repeated hydrochloric acid exchange followed by lyophilization ^64^. For pharmacological inhibition of C5aR1 signaling, we injected the purified peptide (10 mg/kg) subcutaneously four times on alternate days after commencing AQP4-IgG infusion. PBS was injected into a sham-treated group.

### Nissl staining and analysis

Lumbar cord sections were rehydrated in 1× PBS (pH 7.2) for 30 min, followed by permeabilization in 1× PBS containing 0.1% Triton X-100 for 10 min. Sections were then washed twice (5 min each) in 1× PBS. NeuroTrace staining (1:150 dilution in 1× PBS) was applied at room temperature for 20 min in a 96-well plate. After staining, sections were washed in 1× PBS containing 0.1% Triton X-100 for 10 min, followed by two additional 5-minute washes in 1× PBS at room temperature. Sections were washed overnight at 4, mounted and coverslipped using SlowFade™ Diamond Mountant (Thermo Fisher Scientific), and stored in the dark at room temperature to set the mounting medium. Images were acquired as Z-stack planes (3×4 tiling, 1,024 × 1,024-pixel resolution, 2.5 μm Z-steps, 4 slices) using a Zeiss LSM980 Airyscan2 confocal microscope with ×10 objectives. Nissl count analysis in the lumbar ventral gray matter was performed using maximum-intensity confocal Z-stack images. A uniform threshold was applied to define the Nissl signal ROI. The number of Nissl-positive cells and Nissl signal area were quantified using the ImageJ “Analyze Particle” function (NIH, Bethesda, MD). The % Nissl area was calculated by normalizing total Nissl signal area to the ventral gray matter outline.

### Lipid-droplet (LD) staining and analyses

LD staining was slightly modified from a previous description ^44^. Post-fixed lumbar cord sections were rinsed in 1× PBS for 5 min and incubated in 1× PBS containing 0.25% Triton X-100 before 10-minute incubation with BODIPY 493/503 (Thermo Fisher, D3922, 1 μg/mL). To prevent dye washout, sections were transferred directly from the staining solution to glass slides without additional washing. To image LDs, we used a Zeiss LSM980 confocal microscope at ×63 magnification (2,048 × 2,048 pixels, 0.5 μm Z-steps), keeping constant the Z-slice numbers per lumbar section within each experiment. Unbiased quantification was conducted using the “Spot model” for LD counts and the “Surface model” for representative images in Imaris v10.0.

### Mouse astrocyte culture and immunostaining

*AQP4^−/−^* mice were created and bred in-house (Lennon laboratory, Mayo Clinic). Mixed glial cultures were established from newborn mouse cerebral cortices as previously reported ^65^ and used at weeks 2-5. Microglia were depleted by adding Clodrosome (100 μg/mL; Encapsula Nano Sciences LLC) to growth medium (DMEM, 10% FCS, sodium pyruvate, glutamine, antibiotics) for 72 hrs from 2-5 weeks of initial plating and was confirmed using guinea pig anti-IBA1 IgG immunostaining (Synaptic Systems; No. 234 308). For immunofluorescence experiments, astrocytes were plated on glass coverslips with a uniform application of poly-D-lysine and laminin (Corning BioCoat; 354087), postfixed in 4% PFA for 15 min, washed, permeabilized 5 min in PBS/0.02% Trion X-100 and blocked in 3% donkey serum, 30 min. Cells were incubated in primary antibodies at 4 for 16 hrs, washed in 1× PBS, and exposed to secondary antibodies, mounted with SlowFade™ Diamond Antifade Mountants (Thermo Fisher Scientific), and imaged 1 frame by Zeiss 980 microscope. Primary antibodies: mouse anti-GFAP (Abcam; ab4648, 1:1,000), rabbit anti-AQP4 (Sigma; A5971, 1:500), guinea pig anti-IBA1 (Synaptic Systems), rat anti-CXCL1 (R & D, 1:200). Alexa fluor-conjugated secondary antibodies (donkey anti-mouse IgG, 647, donkey anti-rabbit IgG, 488, donkey anti-guinea pig Cy3, and donkey anti-rat IgG, 594) were diluted 1:500 in PBS without permeabilization for 2 hrs at RT. Data represent neurons (*n*) from multiple independent experiments.

### Neutrophil and microglia ablation

To deplete peripheral neutrophils, mice were injected twice *i.p.* with anti-mouse Ly6G (*InVivo*MAb, BioXCell, BE0075-1, Clone: 1A8, 100 mg/kg) or isotype control rat IgG2a, at 5 and 3 days before rotarod testing. The following antibodies identified neutrophils (Gr1 CD11b, Gr1 MPO□) in peripheral blood using spectral flow cytometry (Cytek Aurora, Cytek Biosciences): CD45-V450 (Tonbo, 75-0451-U100), CD45-PE-CF594 (BD, 562420, 1:1,000), CD11b-PE-Cy5 (Tonbo, 55-0112-U100, 1:1,000), Ly6G-APC-Cy7 (BioLegend, 127623, 1:200), Ly6C-PerCP (BioLegend, 128028, 1:100), Gr1-PE (BioLegend, 108408, 1:200), MPO (R & D, AF3667, 1:1,000), Alexa Fluor 488-conjugated (anti-rat, Invitrogen, A-21208).

For microglia ablation, mice were fed *ad libitum* chow containing PLX3397 [colony-stimulating factor 1 receptor (CSF1R) inhibitor, 600 mg/kg, Chemgood, C-1271] for at least 3 weeks prior to surgery and behavioral tests.

### Data analysis and statistics

All statistical analyses were conducted by GraphPad Prism v8 (GraphPad Software Inc.). Normality and variance were tested using the Shapiro-Wilk test. For comparisons between two groups, an unpaired Student *t*-test was performed. When comparing three groups, one-way ANOVA followed by Tukey *post hoc* test was used. For comparisons involving four groups with two independent variables (e.g., treatment × genotype), two-way repeated measures ANOVA with Sidak *post hoc* test were applied, including in experiments such as *C5aR1* deficiency validation. Additionally, for repeated measures, two-way repeated-measures ANOVA with Sidak post hoc test was used, as in the Rotarod test. Sample sizes were determined based on previous studies utilizing similar methodologies. Data are presented as mean ± standard error of the mean (SEM). A *p*-value < 0.05 was considered statistically significant; non-significant results (*p* > 0.05) are not displayed.

### List of Supplementary Materials

Materials and Methods

Figure S1 to S9 for multiple supplementary figures

Movies S1 to S5

References (*1*–*65*)

## Acknowledgments

We thank Olivia Knopke-Mooney for technical assistance, Dr. Na Wang for technical advice, Dr. Trace A. Christensen (Mayo Microscopy and Cell Analysis Core) and Dr. Koichiro Haruwaka for acquisition and 3D rendering of the serial electron-microscopy, and Dr. Jiaying Zheng, Mark A. Maynes and Aubrey Liew for advice and assistance with spectrum flow cytometry. The graphic abstract and cartoon were created by Bio Render (https://www.biorender.com/).

## Funding

National Institutes of Health grants R01 NS110949 (L-JW, VAL) and R35 NS132326 (L-JW).

## Author contributions

Conceptualization: FQ, L-JW, VAL

Methodology: FQ, SZ, T-JC, YG, CL

Investigation: FQ, SZ, HD

Data Acquisition and interpretation: FQ, YG

Funding acquisition: L-JW, VAL

Project administration: VAL, L-JW

Supervision: VAL, L-JW, CL

Writing – original draft: FQ

Writing – review & editing: FQ, L-JW, VAL

## Competing interests

VAL receives royalties from a patent held by Mayo Clinic Foundation for clinical serological testing for AQP4 autoantibodies and from sale of mouse monoclonal anti-AQP4-IgGs (ECD, AQP4-m21, and CCD, AQP4-m5) licensed to Sigma-Aldrich as MABN2471 and MABN2470.

## Data and materials availability

All data associated with this study will be provided by the lead contact upon reasonable request.

## Supplementary Figure Legends

**Figure S1.**
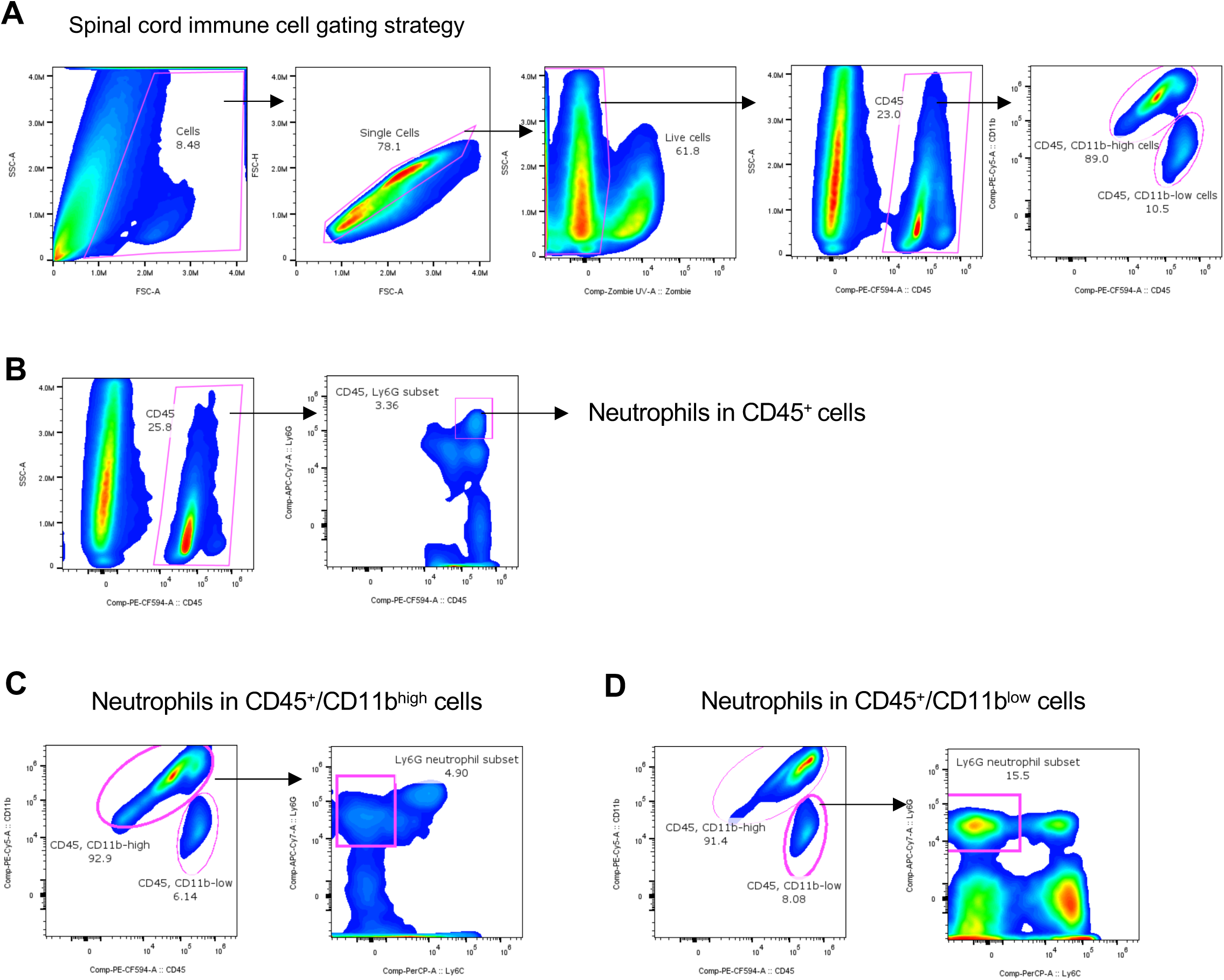
Gating strategies to identify neutrophils in spinal cord by Cytek Aurora system. Related to Figure 1 and Figure 4. Flow cytometry gating strategies to investigate the frequency of Ly6G^+^ neutrophils among CD45^+^/CD11b^+^ immune cells enzymatically dissociated from spinal cord tissues of indicated mice after transcardiac washout of vasculature.

**Figure S2.**
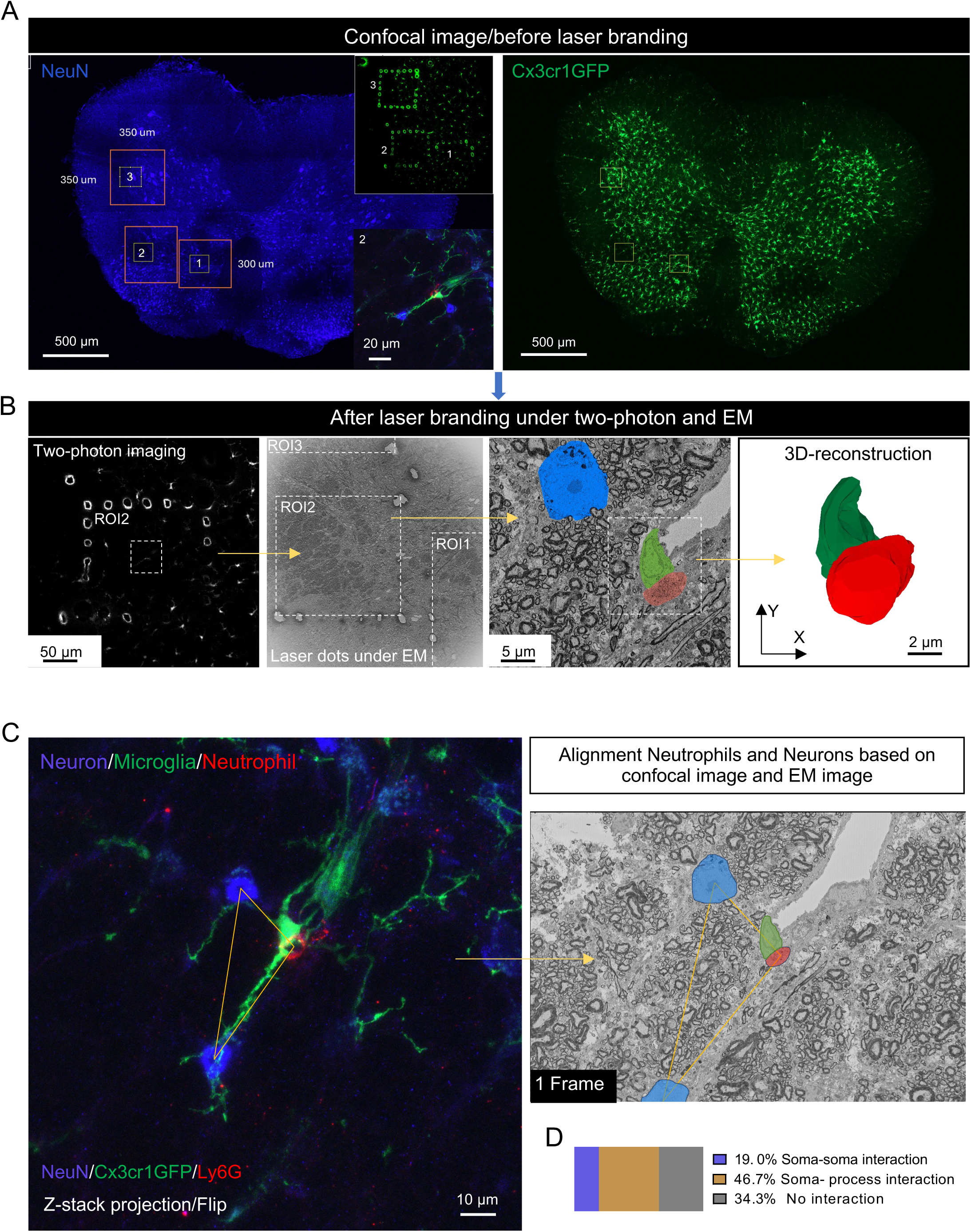
Near-infrared branding by two-photon laser aligns confocal and electron microscopic images to confirm microglia-neutrophil interaction. Related to Figure 2. A. Regions of interest (ROI) were identified in a whole mount lumbar section by confocal microscopy before laser branding. NeuN^+^ neurons (left) and Cx3cr1-GFP^+^ microglia (right) were imaged. B. After laser branding, under 2-photon imaging, the region ROI2 (outlined by branding sites) was relocated by electron microscopy. The same interacting microglial-neutrophil pair was imaged by high resolution electron microscopy and subjected to 3D-serial reconstruction. C. Alignment of neutrophils and neurons according to the confocal Z-stack images and electron microscopy images. The triangular pattern connecting cell somata indicated good alignment.

**Figure S3.**
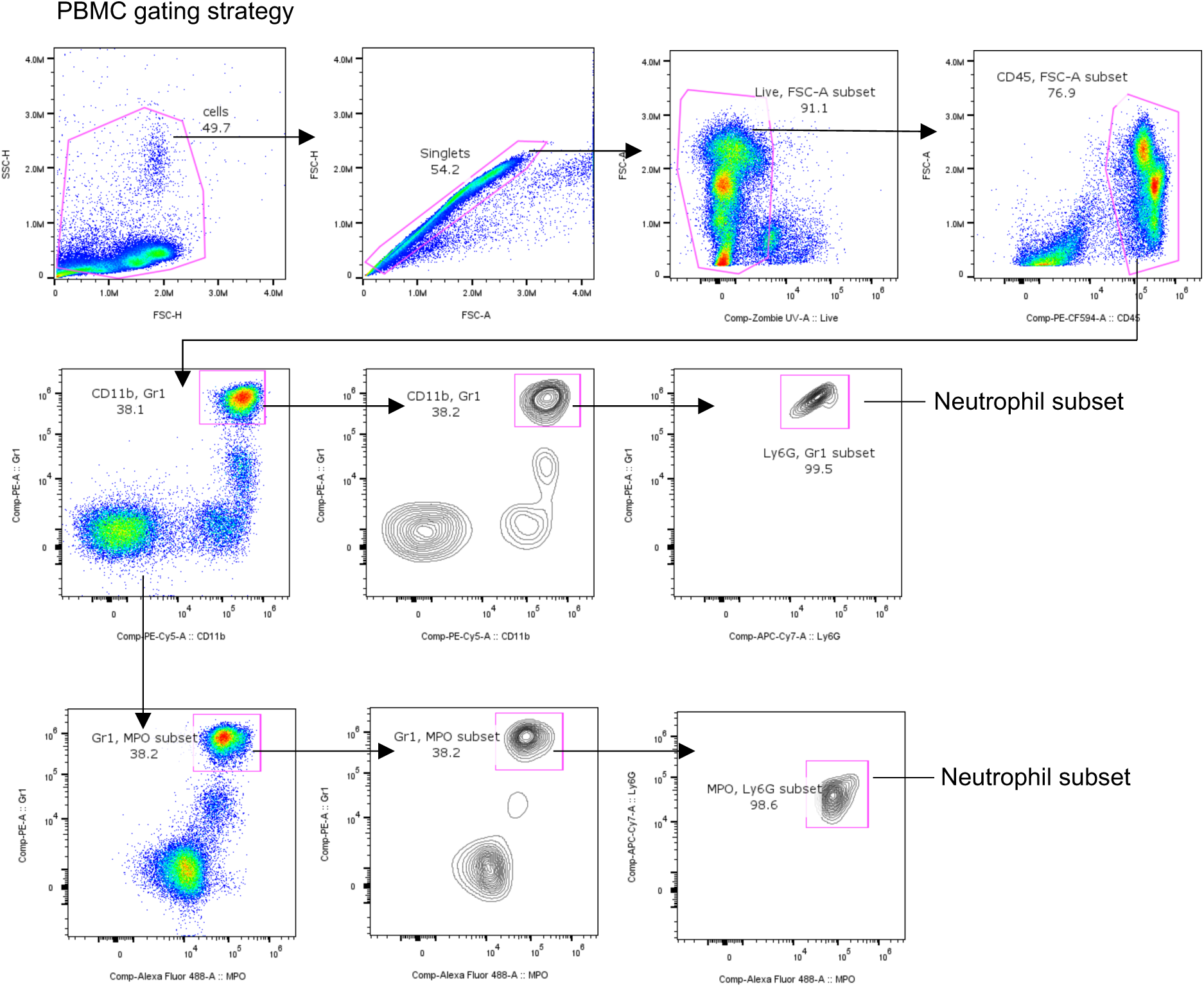
Gating strategies to assess neutrophil ablation efficiency in PBMC. Related to Figure 3. Flow cytometry gating strategies to determine the frequency of Ly6G^+^Gr1^+^ and Ly6G^+^MPO^+^ neutrophil subsets among peripheral blood mononuclear cells (PBMCs).

**Figure S4.**
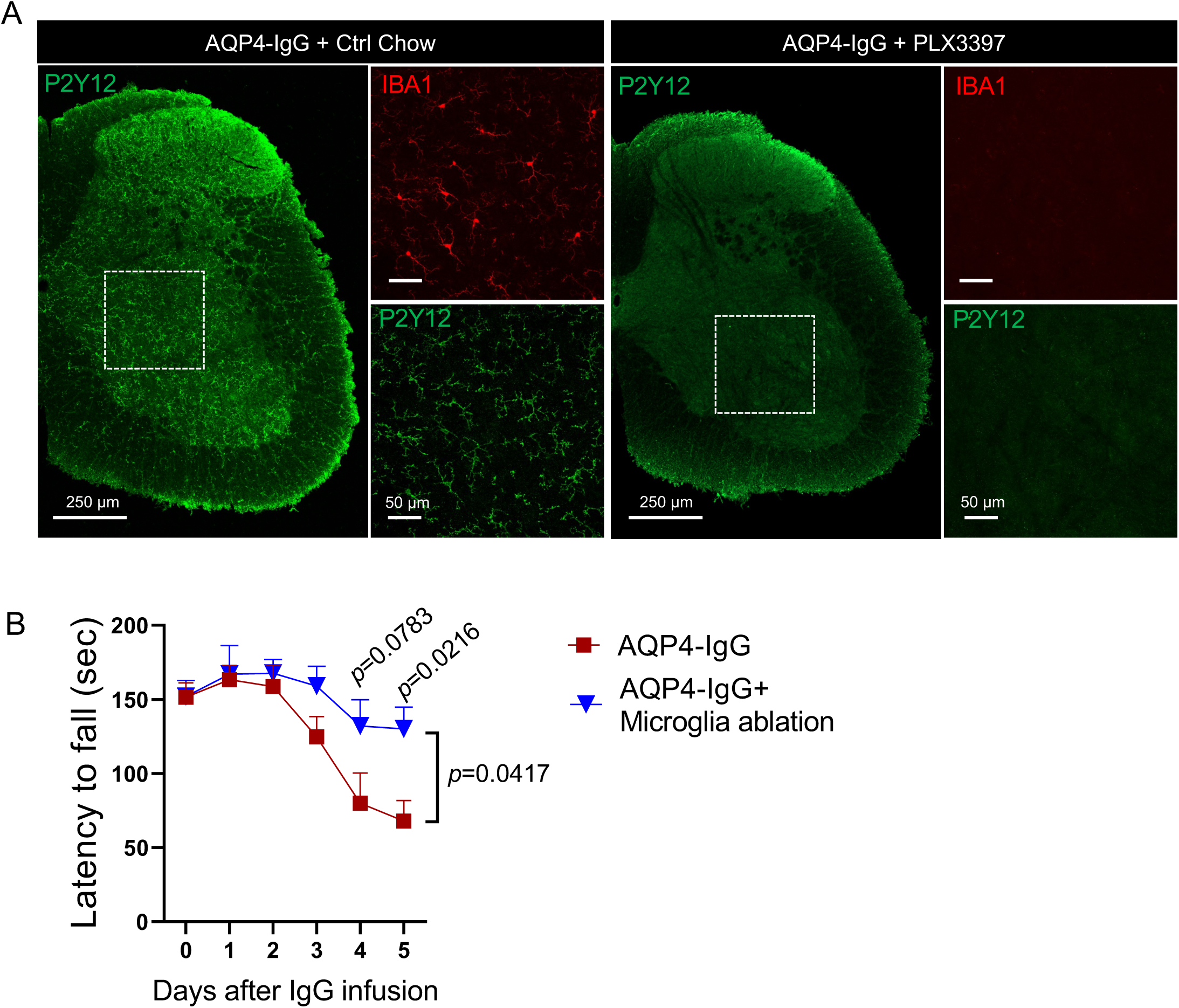
Motor impairment was ameliorated by microglial ablation in NMO mouse model. A. Representative images of P2Y12^+^ (green) and IBA1^+^ (red) microglia in L4 spinal cord sections of mice infused for 5 days with AQP4-IgG after continuous feeding of control chow or microglia-depleting PLX3397 chow starting 24 days before subarachnoid infusion. B. Data for motor impairment, assessed by Rotarod testing, were analyzed by two-way (time × treatment) repeated ANOVA following Sidak multiple comparisons test using GraphPad Prism 8. *F* _(1, 8)_ = 5.865, *p*=0.0417 for treatment (microglial ablation or non-ablation); *F* _(5, 40)_ = 9.459, *p* < 0.0001 for time; *n* = 5 mice per group.

**Figure S5.**
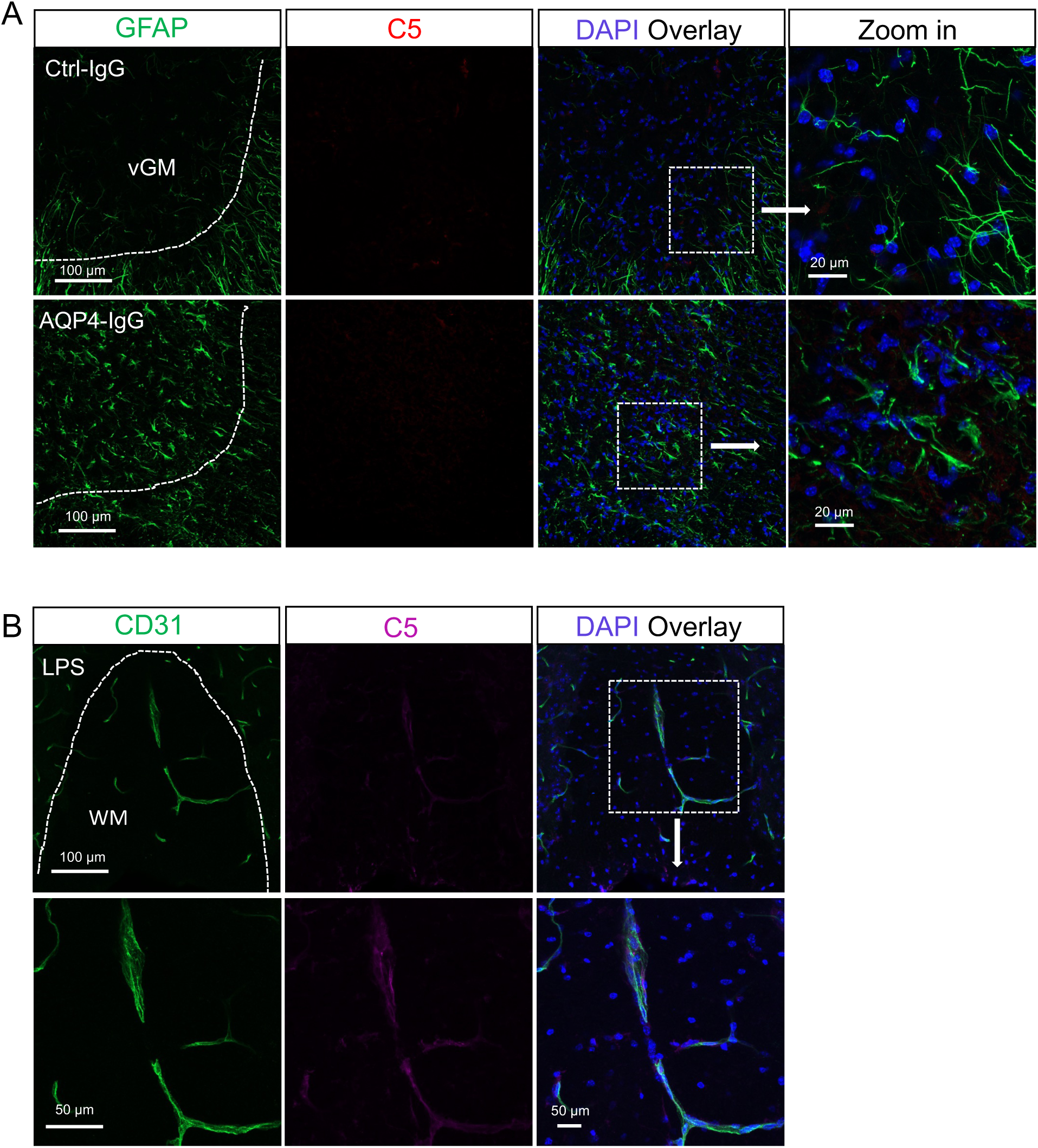
Astrocytes do not express C5 proenzyme in NMO mouse model. Related to Figure 4. A. No complement C5 immunoreactivity was detected in the cytoplasm of astrocytes (GFAP^+^) or any other parenchymal resident or infiltrating cells in lumbar cord of mice infused with either Control-IgG or AQP4-IgG. vGM: ventral grey matter. B. Positive control image demonstrates C5-immunoreactivity within the lumen of a penetrating CD31^+^ blood vessel in spinal white matter of a WT mouse injected i.v. with LPS (1.0 mg/kg, 50 µL).

**Figure S6.**
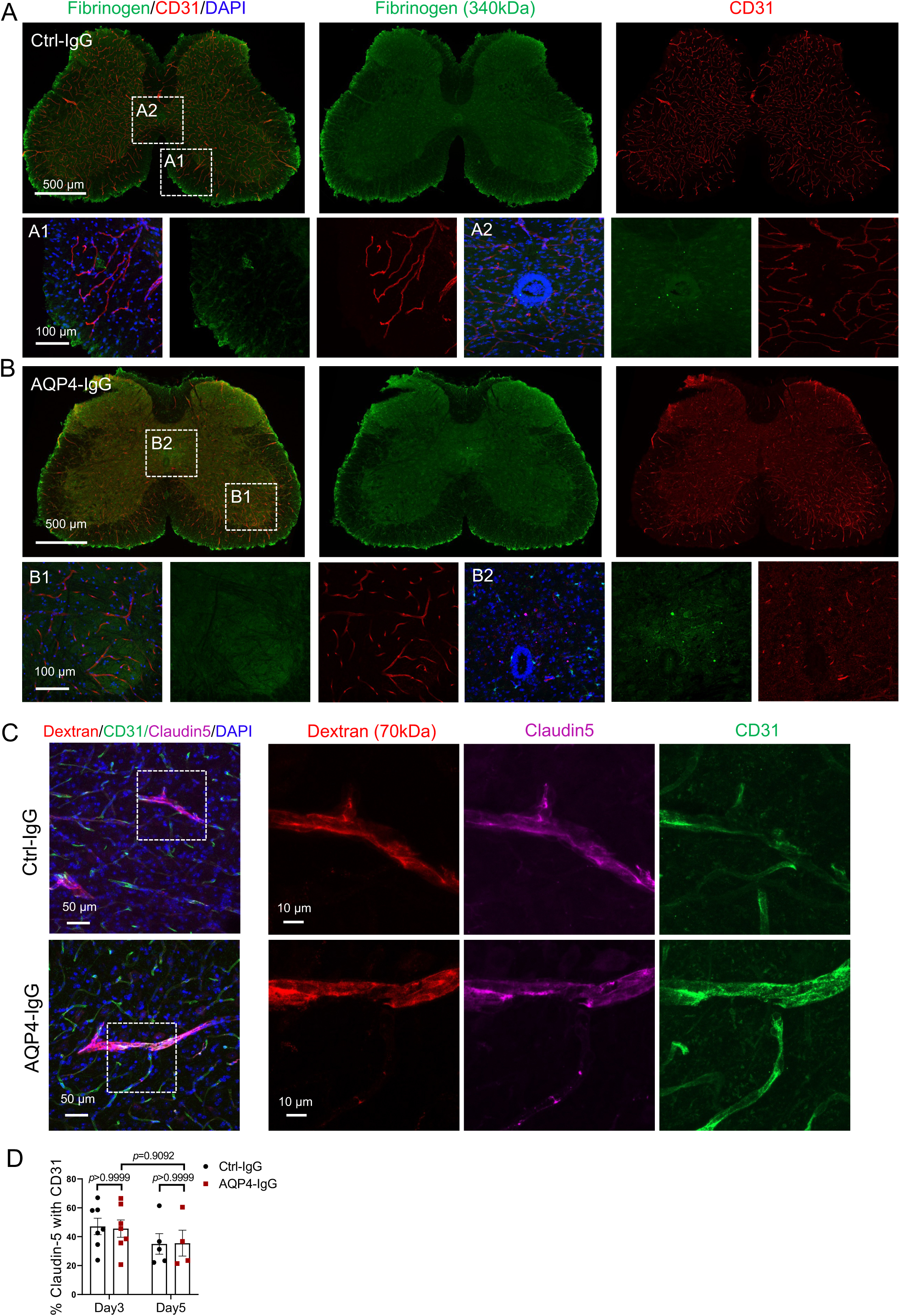
Blood-brain barrier integrity is preserved at day 3 in this NMO mouse model. A-B. Representative images of blood vessel (CD31^+^, red) and fibrinogen (green) immunoreactivities in lumbar cord sections from Control-IgG recipient mice (A) and AQP4-IgG recipient mice (B). Merged images, left. Boxed areas from Control-IgG recipient mice are enlarged in A1 and A2. Boxed areas from AQP4-IgG recipients are enlarged in B1 and B2. C. Confocal images of blood vessel (CD31^+^, green), Dextran red (70 kDa) and Claudin 5 (green) immunoreactivities in lumbar cord sections from Control-IgG recipient mice and AQP4-IgG recipient mice. D. Quantification of the percentage area occupied by Claudin-5 of CD31 blood vessels in lumbar sections from day 3 and day 5 of AQP4-IgG and Ctrl-IgG infused mice. (Treatment: *F* _(1, 19)_ = 0.0055, *p*=0.9417; time: *F* _(1, 19)_ = 2.600, *p*=0.1233; *n* = 4-7 mice per group).

**Figure S7.**
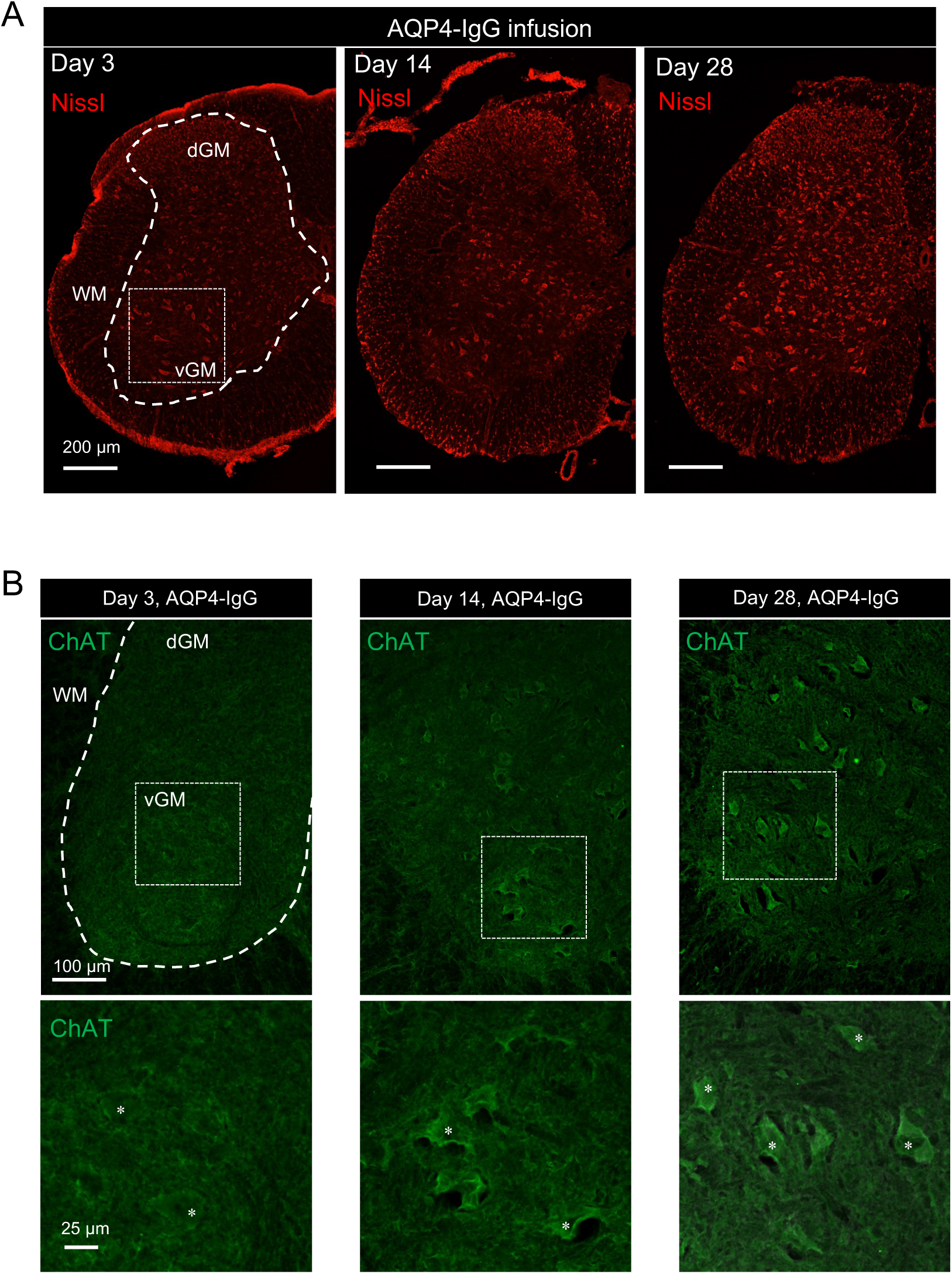
Neuronal dysfunction is reversible in this NMO mouse model. Related to Figure 6 B-D. A. Representative images of neuronal Nissl body staining in L4 cord sections of AQP4-IgG recipient mice at days 3, 14 and 28. AQP4-IgG infusion stopped at day 7. vGM: ventral grey matter; dGM: dorsal grey matter. B. Representative images of ChAT-immunoreactive motor neurons in L4 cord sections of AQP4-IgG recipient mice at days 3, 14 and 28; WM: white matter (upper). Asterisks indicate cytoplasmic ChAT-immunoreactivity, which is less intense at day 3 (lower).

**Figure S8.**
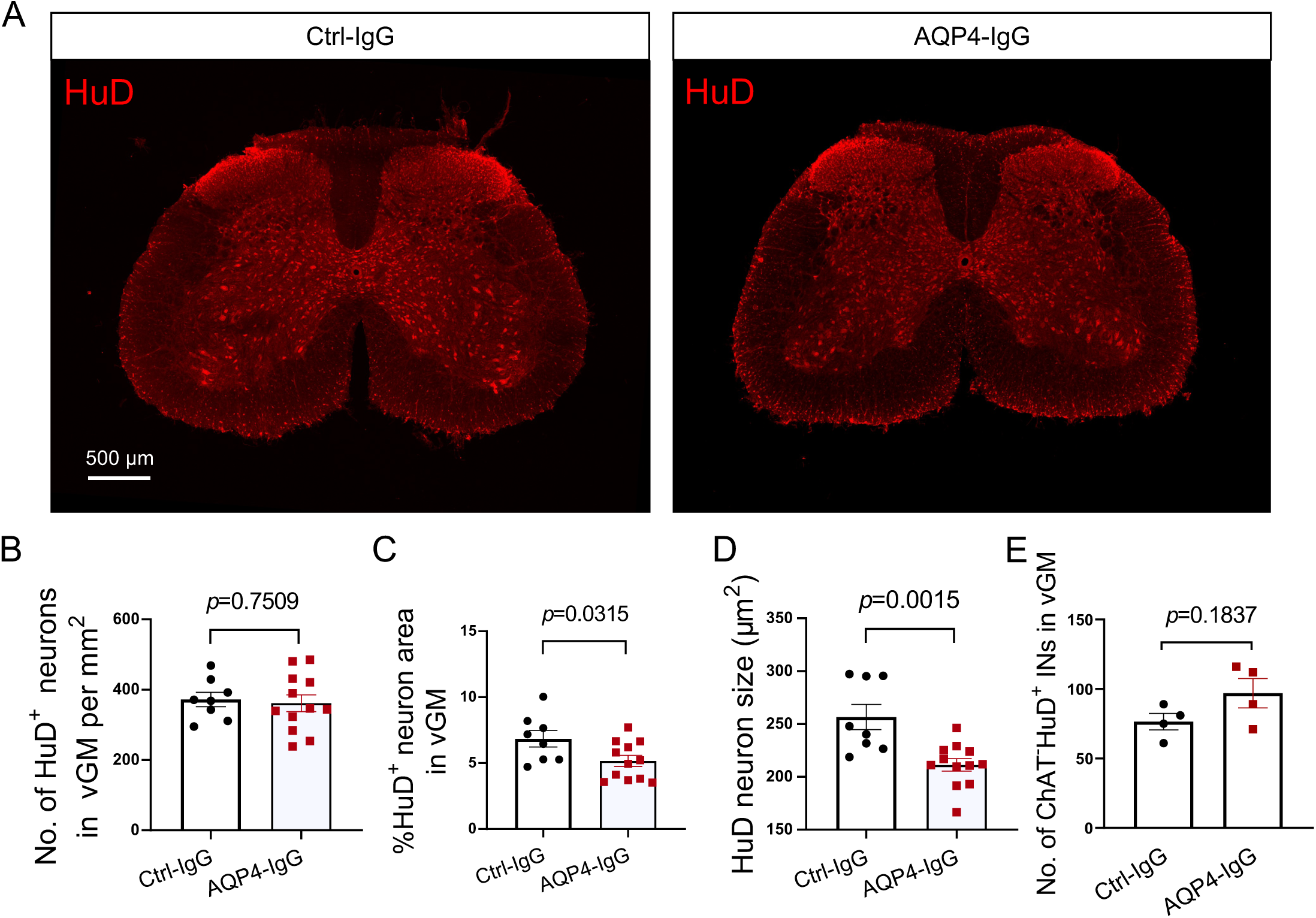
Neuronal changes consistent with functional impairment in this NMO mouse model. A. Confocal images show HuD-immunoreactivity (RNA-binding protein antigen of ANNA1-IgG) in neurons within representative L4 cord sections of mice infused by subarachnoid route with Control-mouse IgG or mouse monoclonal AQP4-IgG. (*n* = 8-12 sections from 4-6 mice per group). B-D. The numbers of HuD^+^ neurons in vGM did not differ significantly (B); the percentage of HuD^+^ neuron area (C) and HuD^+^ neuronal average size (µm^2^) (D) in the vGM of L4 cord are comparable in recipients of Control-IgG and AQP4-IgG. (*n* = 8-12 sections from 4-6 mice per group). vGM: ventral grey matter. E. Quantification of the number of ChAT^-^HuD^+^ interneurons in the vGM of L4 cord. (*n* = 4 mice per group). Data represent means ± SEM and all statistical tests are two-sided. Unpaired Student t-test. *p* < 0.05 was considered a significant difference.

**Figure S9.**
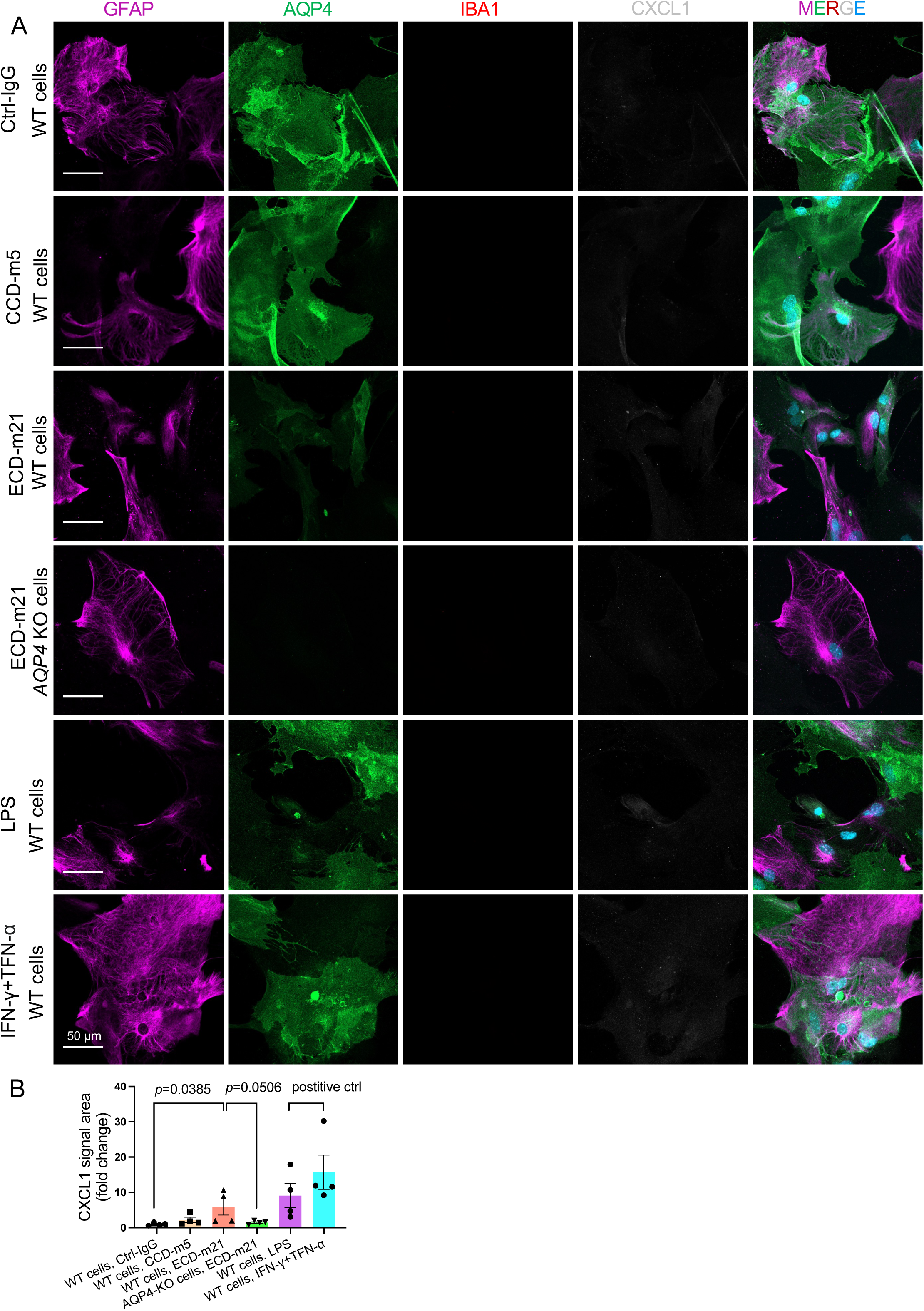
Upregulation of astrocytic CXCL1 production in response to pathogenic AQP4-IgG is greatly diminished in mouse glial cultures depleted of microglia. A. Microglia were depleted from primary cultures of neonatal mouse glia by adding Clodrosome (100 µg/mL; Encapsula Nano Sciences LLC) for 72 hrs. No microglial immunostaining (IBA1) was detected. Astrocytic cytoplasm is identified by GFAP immunostaining; AQP4-immunoreactivity was abundant on astrocytes exposed to control mouse IgG or a non-pathogenic monoclonal mouse IgG specific for the AQP4 cytoplasmic C-terminal domain (CCD; negative control) but was largely cleared by endocytosis-degradation induced by exposure to the pathogenic monoclonal mouse IgG specific for the AQP4 extracellular domain (ECD) and is not expressed on astrocytes derived from AQP4-knock-out mice. B. CXCL1 signal intensity was quantified from A images. Data represent means ± SEM and all statistical tests are two-sided. (*n* = 4 wells). One-way ANOVA with Tukey *post hoc* in A. *p* < 0.05 was considered a significant difference.

**Movie S1.** Tracing of neuron (NeuN^+^, blue), microglia (Cx3cr1GPF^+^, green), and neutrophil (Ly6G^+^, red) based on serial electron microscopic images, related to Figure 2C.

**Movies S2-1, S2-2.** Interacting microglia (Cx3cr1GFP^+^, transparent, S2-1; non-transparent, S2-2) and neutrophil (Ly6G^+^); video created by Animation function in Imaris based on confocal Z-stack images, related to Figure 2E.

**Movie S3.** Interacting microglia (IBA1^+^) and netting neutrophil (MPO^+^); nuclear, blue (DAPI); video created by Animation function in Imaris based on confocal Z-stack images, related to Figure 2F.

**Movie S4.** Behavioral video showing Rotarod motor performance of mice infused with Ctrl-IgG (left) or pathogenic AQP4-IgG (right).

**Movie S5.** Behavioral video showing Rotarod motor performance of mice infused with AQP4-IgG alone (right 2 mice) or combined with anti-CXCL1-IgG (left 3 mice).

